# Color-neutral and reversible tissue transparency enables longitudinal deep-tissue imaging in live mice

**DOI:** 10.1101/2025.02.20.639185

**Authors:** Carl H.C. Keck, Elizabeth L. Schmidt, Richard H. Roth, Brendan M. Floyd, Andy P. Tsai, Hassler B. Garcia, Miao Cui, Xiaoyu Chen, Chonghe Wang, Andrew Park, Su Zhao, Pinyu A. Liao, Kerriann M. Casey, Wencke Reineking, Sa Cai, Ling-Yi Zhang, Qianru Yang, Lei Yuan, Ani Baghdasaryan, Eduardo R. Lopez, Lauren Cooper, Han Cui, Daniel Esquivel, Kenneth Brinson, Xiaoke Chen, Tony Wyss-Coray, Todd P. Coleman, Mark L. Brongersma, Carolyn R. Bertozzi, Gordon X. Wang, Jun B. Ding, Guosong Hong

## Abstract

Light scattering in biological tissue presents a significant challenge for deep *in vivo* imaging. Our previous work demonstrated the ability to achieve optical transparency in live mice using intensely absorbing dye molecules, which created transparency in the red spectrum while blocking shorter-wavelength photons. In this paper, we extend this capability to achieve optical transparency across the entire visible spectrum by employing molecules with strong absorption in the ultraviolet spectrum and sharp absorption edges that rapidly decline upon entering the visible spectrum. This new color-neutral and reversible tissue transparency method enables optical transparency for imaging commonly used fluorophores in the green and yellow spectra. Notably, this approach facilitates tissue transparency for structural and functional imaging of the live mouse brain labeled with yellow fluorescent protein and GCaMP through the scalp and skull. We show that this method enables longitudinal imaging of the same brain regions in awake mice over multiple days during development. Histological analyses of the skin and systemic toxicology studies indicate minimal acute or chronic damage to the skin or body using this approach. This color-neutral and reversible tissue transparency technique opens new opportunities for noninvasive deep-tissue optical imaging, enabling long-term visualization of cellular structures and dynamic activity with high spatiotemporal resolution and chronic tracking capabilities.

**Significance Statement:** Tissue scattering represents a major barrier to deep tissue imaging *in vivo*. We recently showed that tissue can be rendered transparent in the red spectrum using intensely absorbing dye molecules. Here, we introduce a new, color-neutral and reversible tissue transparency approach. We demonstrate longitudinal structural and functional imaging in the deep tissue of awake mice.

## Introduction

Biomedical optics have transformed our understanding of biology through techniques such as widefield and confocal fluorescence microscopy (1), two-photon and three-photon microscopy (2, 3), optical coherence tomography (4), and photoacoustic microscopy (5). A common challenge for all optical imaging approaches is the scattering and absorption of photons in biological tissues, which inherently limits light penetration and imaging depths (6). In most biological tissues, scattering is the primary barrier to deep tissue imaging (7). Scattering arises from the complex structure of biological matter, composed of diverse chemical components with distinct refractive indices (RIs) intermixed at microscopic scales (8). This limitation restricts penetration depth to approximately 100 μm for confocal fluorescence microscopy (9) and no more than 1.5 mm for state-of-the-art multiphoton microscopy using complex and expensive customized equipment (10).

Over the past decades, several methods have been developed to address scattering in biological tissues. By leveraging the inverse relationship between emission wavelength and scattering coefficient, second near-infrared (NIR-II) fluorescence imaging, also known as short-wave infrared (SWIR) fluorescence microscopy, utilizes fluorescent emitters in the NIR-II spectrum (1,000–3,000 nm) to achieve deep-tissue fluorescence imaging in live animals and even humans (6, 11, 12). However, the longer emission wavelengths inherently result in lower spatial resolution due to diffraction limits. Additionally, various optical tissue clearing methods reduce scattering by removing water or lipids from tissues—both essential for sustaining life (13). Consequently, these methods are rarely applied *in vivo*, restricting their utility in dynamic biological systems.

Non-invasive deep-tissue optical imaging in live subjects offers the potential for long-term visualization of cellular structures and dynamic activity, providing high resolution, rapid imaging speed, and chronic tracking capabilities. To this end, our group has recently developed a noninvasive method for achieving tissue transparency without removing any tissue components, offering a potential solution to many of the challenges associated with existing approaches (14). While our technique also reduces RI mismatch by increasing the RI of the aqueous components in tissue, it is fundamentally rooted in the physics of light-matter interaction, employing the Lorentz oscillator model and the Kramers-Kronig relations to analyze various tissue components and strongly absorbing dye molecules (15, 16). This allows RI matching at significantly lower concentrations compared to conventional optical clearing agents, effectively avoiding dehydration and minimizing disruption to biological processes.

Despite the success in achieving optical transparency *in vivo*, our previous work relies on dye molecules with major absorption peaks in the visible spectrum, such as tartrazine, which features peak absorption at 428 nm and lingering absorption up to 600 nm (Fig. 1*A*) (14). Consequently, optical transparency is limited to wavelengths above 600 nm (17), restricting the application of this approach for imaging shorter-wavelength fluorophores, such as commonly used green and yellow fluorescent proteins (GFP and YFP, respectively). In this work, we address this challenge by developing a method to achieve color-neutral and reversible tissue transparency in live mice. By extending skin transparency across the entire visible spectrum, our method enables noninvasive imaging of both YFP and the GFP-based calcium sensor GCaMP in the live mouse brain without removing either the scalp or skull. Additionally, we demonstrate that the creation of a color-neutral transparency window in the mouse scalp allows repeated longitudinal imaging of the same brain regions in the same mouse throughout development. This advancement opens the door to noninvasive deep-tissue optical imaging, providing long-term visualization of cellular structures and dynamic activity with high spatiotemporal resolution and chronic tracking capabilities.

**Fig. 1.**
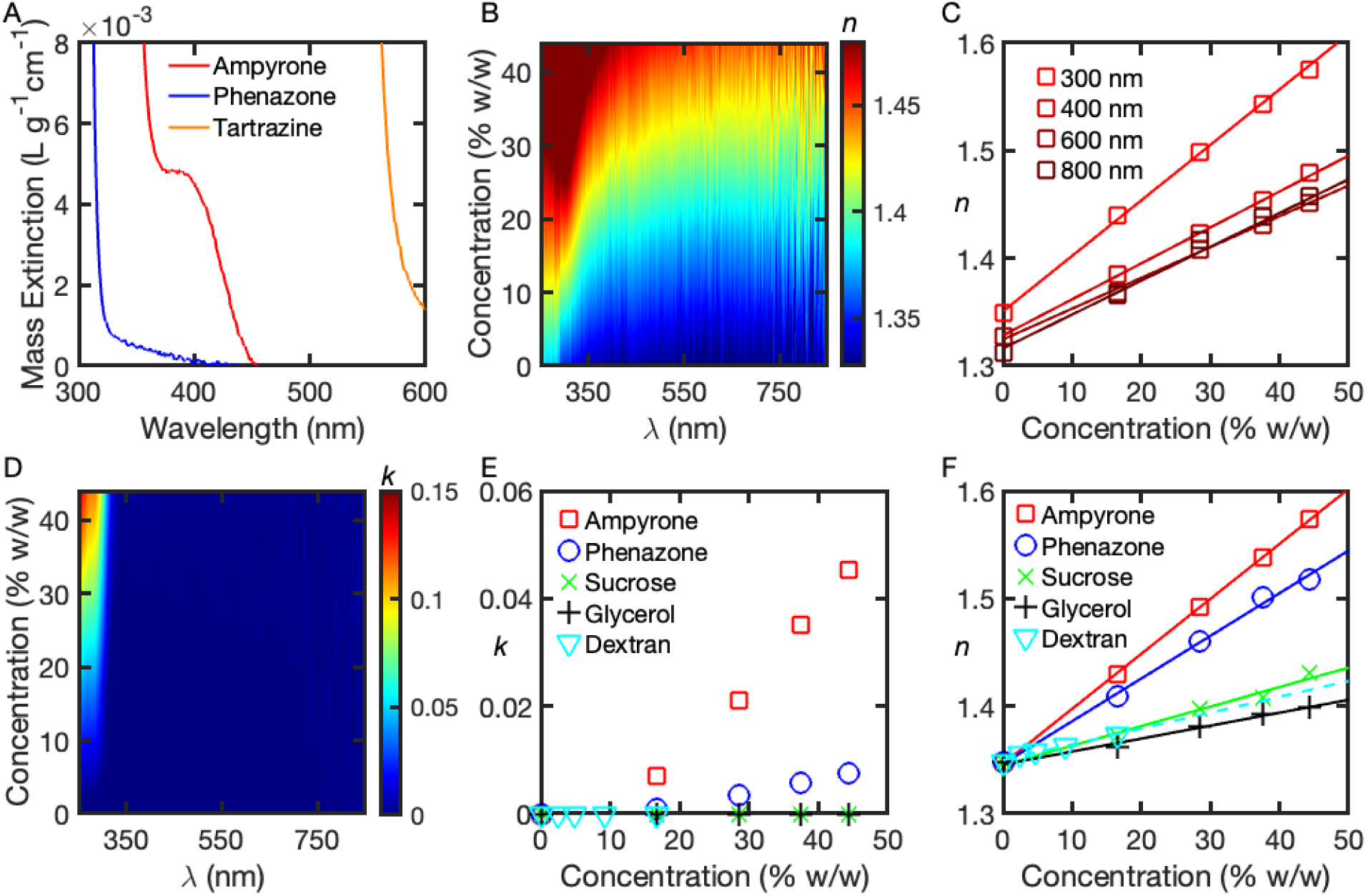
Optical characterizations of ampyrone. (*A*) Comparison of the mass extinction coefficients of phenazone, ampyrone, and tartrazine solutions as a function of wavelength, highlighting the distinct positions of their absorption edges. A 100 mg/mL aqueous solution was used for each spectrum. (*B*) The real RI (*n*) of an aqueous solution of ampyrone as a function of concentration and wavelength. (*C*) The real RI (*n*) of an aqueous solution of ampyrone as a function of concentration at selected wavelengths. (*D*) The imaginary RI (*k*) of an aqueous solution of ampyrone as a function of concentration and wavelength. (*E*) Comparison of *k* for various index-matching agents in aqueous solutions at 310 nm as a function of concentration. (*F*) Comparison of the real RI (*n*) for the same index-matching agents shown in *E* as a function of concentration at 310 nm. The Dextran data had to be linearly extrapolated up to 50% w/w since high concentrations are not achievable due to its solubility limit.

## Results

### Ampyrone is a potent UV absorber that enables color-neutral optical transparency across the visible spectrum

An ideal absorbing molecule should have the majority of its absorption bands in the ultraviolet (UV) spectrum, leaving most of the visible spectrum unobstructed for imaging while leveraging RI modulation in the visible range, enabled by intense UV absorption. This design can exploit the Kramers-Kronig relations, which describe the causal relationship between the wavelength-dependent variation of the real (*n*) and imaginary (*k*) parts of a material’s complex refractive index. These relationships are expressed analytically as follows:

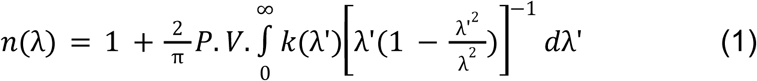

where λ is the wavelength of interest, P.V. is the Cauchy principal value of the integral, *n*(λ) is the real part of the complex RI (referred to hereafter simply as the RI), and *k*(λ) is the imaginary part of the complex RI. The imaginary RI, *k*(λ), is related to the absorption coefficient α(λ) by

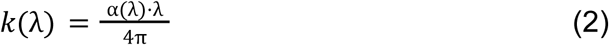

Based on this theoretical framework, an ideal RI-matching agent should maximize integrated absorption below the wavelength of interest while minimizing absorption at the wavelength of interest. This requirement suggests a sharp absorption edge at a wavelength immediately below the desired imaging wavelengths (Fig. S1). Guided by this design principle, we first explored phenazone, also known as antipyrine. Phenazone has been previously reported as an *ex vivo* RI-matching agent (18, 19), due to its high refractive index and low visible absorption. However, effective phenazone solutions typically require concentrations of 55% or higher, resulting in a mixture where phenazone dominates and water molecules constitute less than half of the solution.

We sought to understand the relative inefficiency of phenazone by measuring its UV-Vis transmission spectrum in an aqueous solution. The absorption spectrum of phenazone in the UV-visible range reveals intense absorption below 300 nm, with a rapid decay to minimal absorption beyond 320 nm (Fig. 1*A*). For fluorescence imaging with GFP and more red-shifted fluorophores, there is a significant gap of at least 160 nm (from the sharp absorption edge at 320 nm to GFP’s excitation wavelength at 488 nm) that does not contribute to RI increase in the visible spectrum. To address this inefficiency, strategies aimed at red-shifting the absorption edge to fill this gap in the absorption spectrum would be an effective approach for identifying more efficient RI-matching agents that function at lower and more physiologically tolerable concentrations.

To red-shift the absorption edge, we applied an empirical principle well known to organic chemists for synthesizing donor-acceptor colorants. Specifically, it is established that electron-donating groups, particularly amine groups, conjugated with an extended chromophore system can induce both bathochromic (red-shifting) and hyperchromic (up-shifting) shifts in the absorption peak (20). This effect arises from the extension of the π system through the involvement of nonbonding p-orbitals and the intensification of the molecular dipole. Specifically, we hypothesized that an amine group addition at the C4 site of the central pyrazole backbone of phenazone can significantly red-shift the absorption edge, thus bringing the resonance closer to the visible wavelengths (14, 21). We also hypothesized that this amine group can create a push-pull electron donor-acceptor system, which is also responsible for the redshift (22). The UV-Vis transmission spectrum of the resulting molecule, ampyrone, exhibits a red-shifted absorption edge extending to approximately 400 nm (Fig. 1*A*), significantly increasing the total absorption area. As a result, ampyrone appears pale yellow (Fig. S2*A*) while otherwise remaining largely color-neutral across the visible spectrum.

### The intense UV absorption of ampyrone leads to efficient RI modulation with minimal absorption across the visible spectrum

Ampyrone exhibits a broad absorption peak at 220-320 nm (Fig. S2*B*), with shoulder peak absorption extending well beyond 400 nm (Fig. 1*A*). As a result, after dissolving in water, ampyrone can significantly increase the real RI, *n*, through the Kramers-Kronig relations (Fig. S2*C*). Specifically, the RI modulation of water as a function of ampyrone concentration (% w/w) and wavelength (nm) (Fig. 1*B*) reveals that peak index modulation occurs at ∼310 nm, after which it rapidly plateaus across the visible and near-infrared wavelengths. We further examined how *n* varies with concentration at multiple selected wavelengths (Fig. 1*C*). Near the RI maximum at 300 nm, within the high-dispersion regime, *n* increases more rapidly with concentration. However, moving into the visible spectrum (400–800 nm), the dependence of *n* on concentration remains relatively stable, reinforcing the plateau behavior.

To further corroborate ampyrone’s intense absorption in the UV spectrum and minimal absorption across the visible spectrum, we plotted the imaginary refractive index (*k*) as a function of both concentration and wavelength (Fig. 1*D*). This analysis reveals several key findings. First, ampyrone exhibits the most intense UV absorption (e.g., at 310 nm) at equivalent concentrations compared to commonly used index-matching agents (Fig. 1*E*). This exceptionally strong UV absorption—far exceeding that of other index-matching agents—underpins its RI modulation efficiency in the visible spectrum (Fig. 1*B&C*). Second, beyond the UV spectrum, ampyrone demonstrates minimal absorption across the entire visible range (Fig. 1*D*), enabling imaging with common fluorescent proteins while minimizing attenuation in the transparency window. Third, as a result of its intense UV absorption and negligible visible absorption, ampyrone achieves the most efficient modulation of the real refractive index (*n*) in aqueous solutions, compared to previously reported *in vivo* index-matching agents (Fig. 1*F*). These agents include sucrose (23), glycerol (24, 25), and dextran (40 kDa) (26) with their full real and imaginary RI spectra as a function of concentration provided in Fig. S3. Lastly, ampyrone’s small molecular weight (203 g/mol) provides significant advantages in water solubility and diffusion efficiency in tissue.

### Ampyrone achieves transparency in scattering phantoms

To further evaluate the optical performance of ampyrone in enhancing tissue transparency, we prepared 5-mm-thick scattering phantoms composed of 1% w/w agarose hydrogel with 1 μm silica particles dispersed within (Fig. 2*A*). These phantoms were backlit with a grid pattern using an LED panel to assess light transmission at varying ampyrone concentrations. Without ampyrone, the grid lines were completely obscured due to strong light scattering through the phantom. As the ampyrone concentration increased, the grid lines became progressively more visible, with maximum clarity observed at 38% w/w (corresponding to *n* ∼1.43, closely matching the RI of silica particles). However, at 44% w/w, where the RI exceeded the optimal matching condition, the clarity of the grid pattern decreased. This decreased clarity confirms that the observed transparency arises from the RI modulation of the aqueous component in the scattering phantom by ampyrone.

**Fig. 2.**
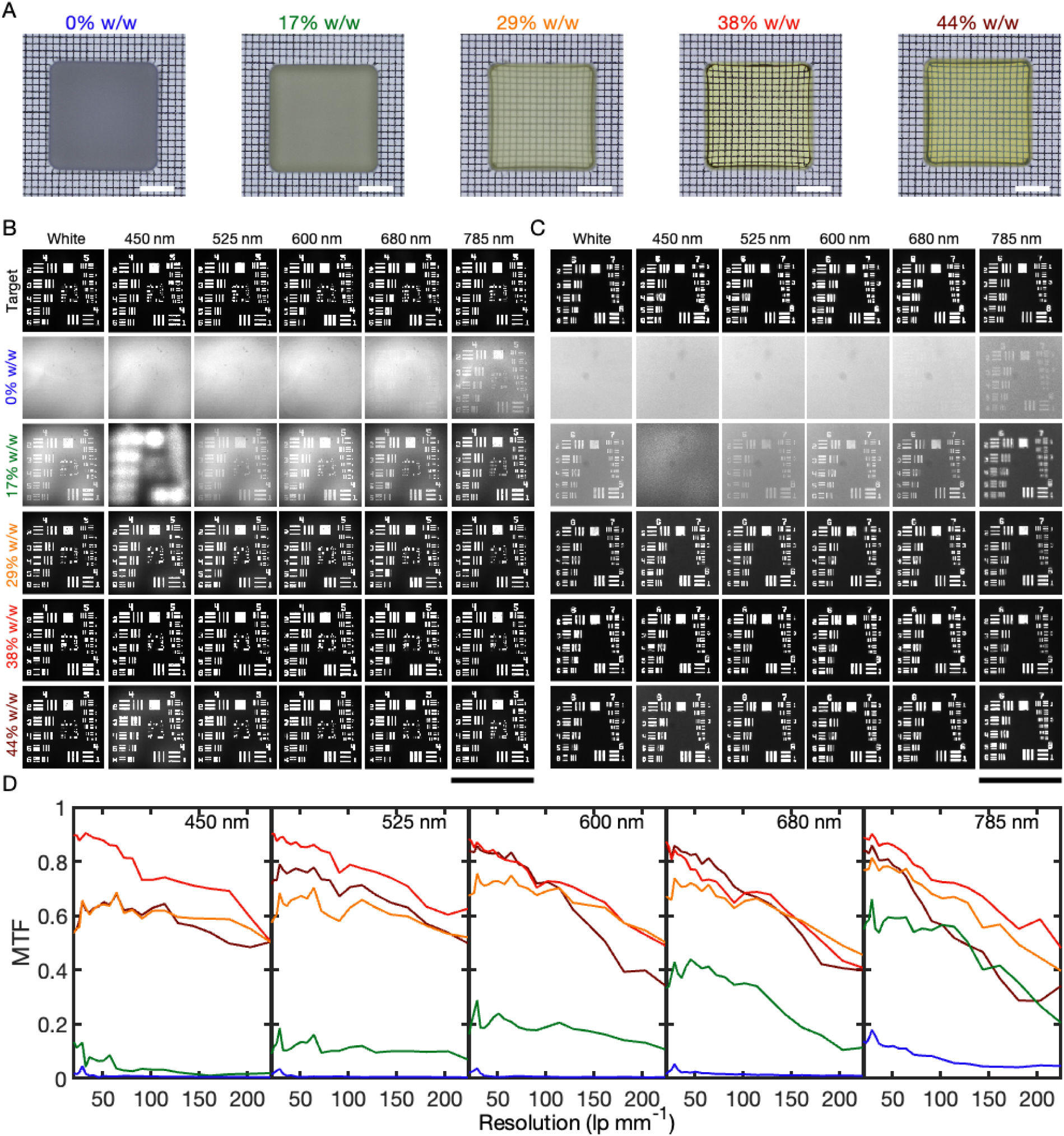
Resolution characterization of scattering phantoms. (*A*) Phantoms of 1 μm silica beads in 5-mm-thick scattering phantoms composed of agarose hydrogel with increasing concentrations of ampyrone. Scale bars are 5 mm. (*B*) 1951 USAF resolution test target images through 1 μm silica beads in 2-mm-thick scattering phantoms. The scale bar is 1.2 mm. (*C*) Same test target images as in *B*, but zoomed in to display the smallest features. The scale bar is 296 μm. (*D*) Modulation transfer functions (MTFs) for all concentrations of ampyrone used in *B* and *C* with the same color coding by concentration. Each subplot shows the MTFs at different concentrations for a single wavelength.

To systematically quantify the relationship between ampyrone concentration and transparency performance in the silica phantom, we prepared phantoms with varying ampyrone concentrations and imaged the transmitted image of a 1951 USAF resolution test target through 2 mm thick phantoms (Fig. 2*B&C*). At 0% w/w (without ampyrone), the USAF target is almost completely obscured by scattering, with only large features visible beyond 785 nm in the near-infrared (NIR) spectrum. As the ampyrone concentration increases, we observe a progressive reduction in scattering and improved image formation across the visible spectrum. At 17% w/w—an undermatched condition—scattering is already noticeably reduced. By 38% w/w, the USAF targets appear nearly identical to the control target without any scattering phantom.

To quantify the resolution improvements due to index matching, we calculated the modulation transfer function (MTF) for each image in Fig. 2B&C. The MTF for all concentrations at a given wavelength is plotted, demonstrating that 38% w/w ampyrone is optimal for increasing the MTF of the scattering phantom across both the visible and NIR spectra. Even at lower concentrations, there is a significant improvement in MTF compared to the original scattering phantom, suggesting that if perfect RI matching is not required, lower ampyrone concentrations can still yield an observable transparency effect.

### Ampyrone achieves optical transparency in *ex vivo* skin

We next evaluated ampyrone’s ability to enhance optical transparency in biological tissue *ex vivo*. To do this, we harvested abdominal skin from 4-month-old mice and incubated it in ampyrone solutions of varying concentrations to assess its ability to induce optical transparency. While 17% w/w ampyrone had minimal effect, partial transparency was observed at 29% w/w, and nearly complete transparency was achieved at 38% w/w. However, higher concentrations, such as 44% w/w, resulted in poor transparency (Fig. 3*A*).

**Fig. 3.**
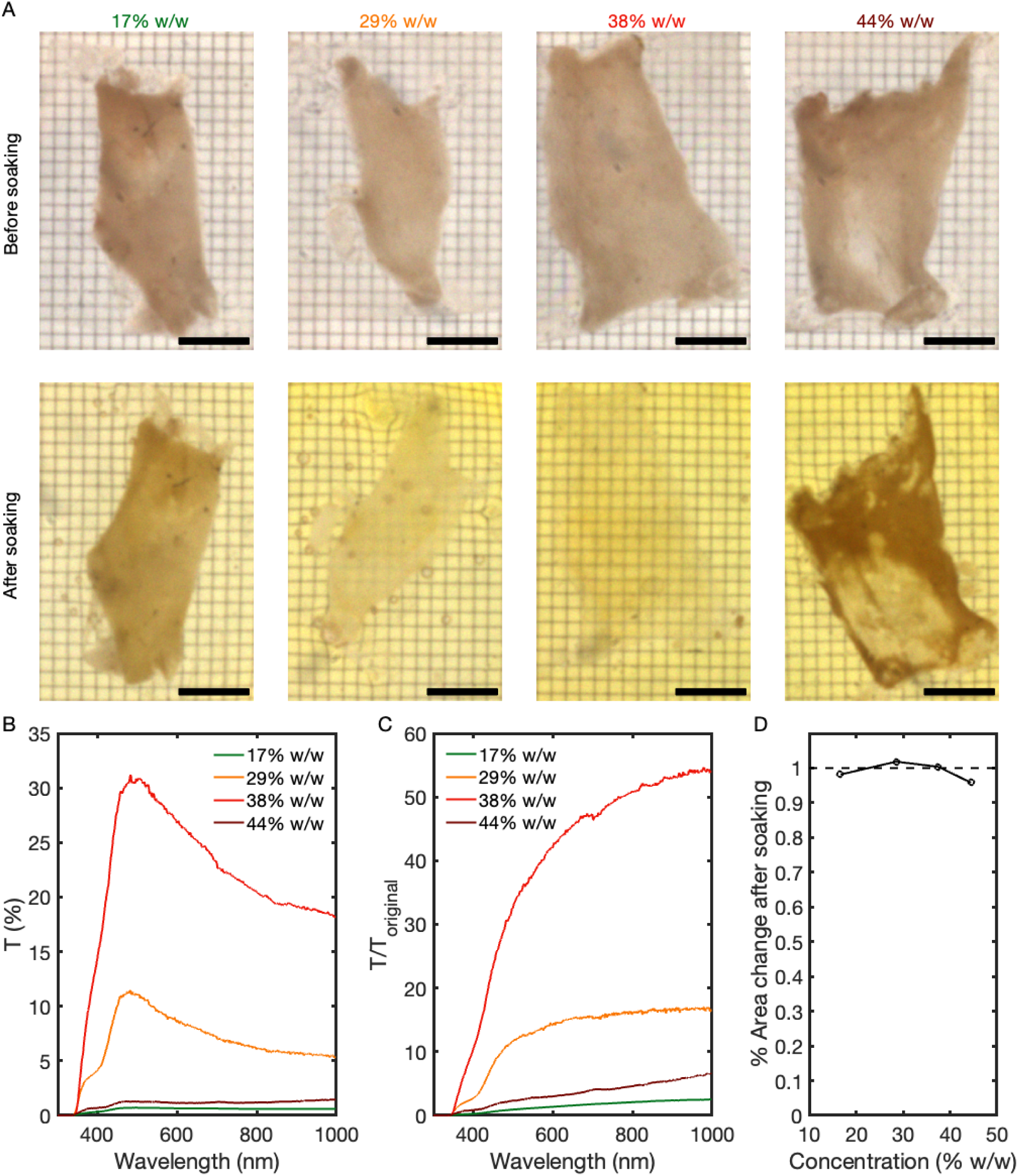
Achieving optical transparency in *ex vivo* mouse skin. (*A*) Mouse abdominal skin before (top) and after (bottom) soaking in ampyrone solutions at various concentrations. (*B)* Transmission spectra through the same four mouse skin samples as in *A* after soaking. (*C*) The ratio of the transmission after soaking to before soaking for the same four mouse skin samples. (*D*) The area change after soaking for the same four skin samples is shown as a function of concentration. All scale bars are 5 mm.

Transmission spectra analysis showed that skin treated with 38% w/w ampyrone reached up to 30% transmission across the visible spectrum, a substantial increase from the <1% transmission of untreated skin (Fig. 3*B*). Quantitative analysis revealed a >50-fold enhancement in transmission for 38% w/w ampyrone compared to untreated samples (Fig. 3*C*). Finally, analysis of skin area before and after treatment showed negligible changes across all tested concentrations, indicating that tissue size was preserved after the tissue became transparent (Fig. 3*D*).

### Ampyrone achieves optical transparency in the live mouse abdomen

Next, we aimed to demonstrate the transparency effect in live mice using a 38% w/w ampyrone solution, which had shown maximum transparency in scattering phantoms and *ex vivo* skin. Although a mouse’s depilated abdominal skin typically appears opaque (Fig. 4*A*), topical application of 38% w/w ampyrone significantly increased transparency, allowing detailed visualization of internal organs such as the liver and intestines within the abdominal cavity (Fig. 4*B*). We confirmed that these emerging features originated from internal organs by physically removing the abdominal skin, which revealed similar organ structures (Fig. 4*C*).

**Fig. 4.**
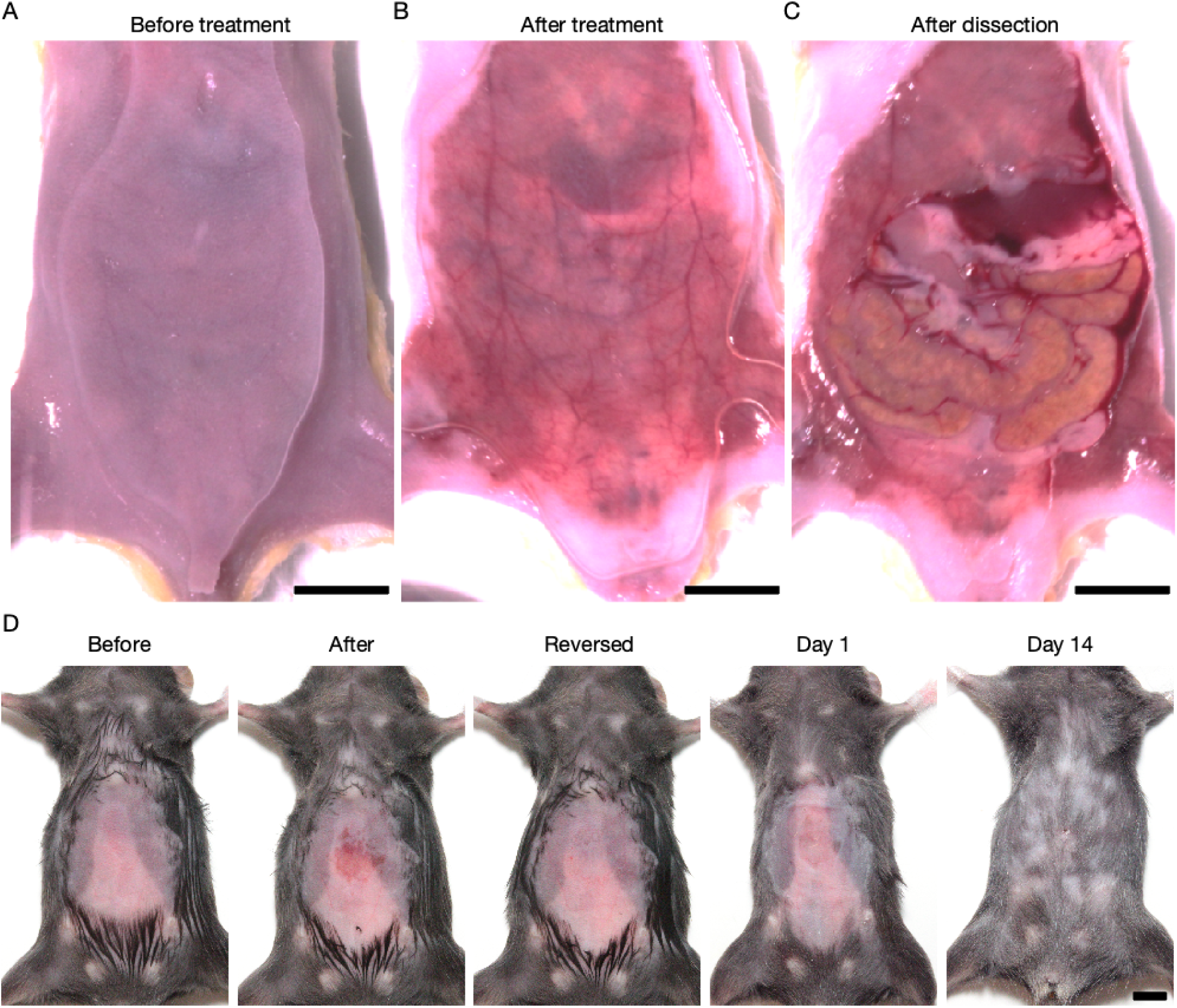
Achieving optical transparency in the live mouse abdomen. (*A*) Mouse abdomen before treatment with ampyrone solution. (*B*) Mouse abdomen shown after treatment with ampyrone solution. (C) The same mouse abdomen in *A*&*B* after dissection. Scale bars are 1 cm in *A*-*C*. (*D*) Images showing the abdominal area of the same mouse before treatment, after achieving a transparent window with ampyrone, after reversing the transparency effect, and hair regrowth on the subsequent days. Scale bar is 1 cm in *D*.

Next, we aimed to demonstrate the reversibility of the ampyrone treatment process (Fig. 4*D*). After creating a transparent window in the abdominal skin using ampyrone, we topically applied a saline solution to extract the ampyrone molecules, effectively reversing the transparency effect and restoring the skin’s original appearance. Following the reversal, we applied a bioadhesive hydrogel to maintain skin hydration before the mouse was recovered from anesthesia (see **Materials and Methods**). The animal exhibited normal activity and behavior in the following days, with the skin appearing normal over the next seven days. Hair regrowth was observed in the treated abdominal area thereafter.

To assess the biosafety of topically applied ampyrone, we conducted blood chemistry and hematology assays on mice that received abdominal treatment with ampyrone. While ampyrone functions as an anti-inflammatory, analgesic, and antipyretic agent (27), it is not entirely biorthogonal as an index-matching agent. The assay results (Fig. S4 & Table S1) showed no statistically significant differences between saline-treated and ampyrone-treated mice at both 1 day and 14 days post-treatment. Histological analysis of skin tissue from mice treated with saline or ampyrone at these time points revealed no significant differences in tissue morphology or inflammatory response (Fig. S5 & Table S2). To further illustrate the minimal impact of ampyrone on live animals, we present a video (Movie S1) demonstrating normal ambulation in an awake mouse immediately following scalp treatment with ampyrone.

We attribute ampyrone’s minimal toxicity to the low dose and transient nature of the treatment. Specifically, we estimate that 20 µL of a 38% w/w ampyrone solution penetrated the skin to achieve transparency, while the subsequent reversal step extracted 80% of the skin-absorbed molecules (see **Materials and Methods**). As a result, only a 0.05–0.17 g/kg body weight dose was effectively administered across all tested animals, significantly lower than the reported LD50 for intraperitoneal ampyrone administration (1.2 g/kg body weight in rodents (27)). This reference on LD50 of ampyrone also highlights ampyrone’s low toxicity as a key factor in its use as a safer analgesic and antipyretic alternative to its parent compound, aminopyrine.

### Ampyrone enables longitudinal two-photon microscopy through the scalp in live mice

Following these demonstrations, we sought to evaluate ampyrone’s potential to enable deep-tissue imaging across the entire visible spectrum, leveraging its color neutrality. We first performed structural imaging of YFP-expressing neurons in the cortex of live Thy1-YFP-H mice through the transparent scalp using two-photon excitation fluorescence microscopy. YFP imaging was not compatible with our previous *in vivo* transparency approach due to the absorption profile of tartrazine, which significantly attenuated YFP fluorescence (14).

Traditionally, two-photon fluorescence microscopy of the mouse cortex requires scalp removal and the installation of a cranial window to replace part of the skull, as these overlying tissues cause significant scattering that prevents microscopic imaging of underlying neurons. This limitation is evident in a three-dimensional (3D) reconstruction of two-photon fluorescence collected to a depth of 700 μm below the depilated scalp surface, where the only observable signal arises from hair follicles within the most superficial 100 μm of the scalp (Fig. 5*A*). In contrast, after treating the scalp with 38% w/w ampyrone to achieve transparency, 3D-reconstructed two-photon fluorescence images revealed extensive Thy1-YFP-H neuronal structures extending up to 700 μm below the scalp surface (Fig. 5*B*). In these images, the scalp, skull, and cortex are distinctly delineated based on their fluorescence properties: hair follicle autofluorescence marks the scalp, an absence of signal (dark layer) represents the skull, and YFP signals highlight cortical neurons.

**Fig. 5.**
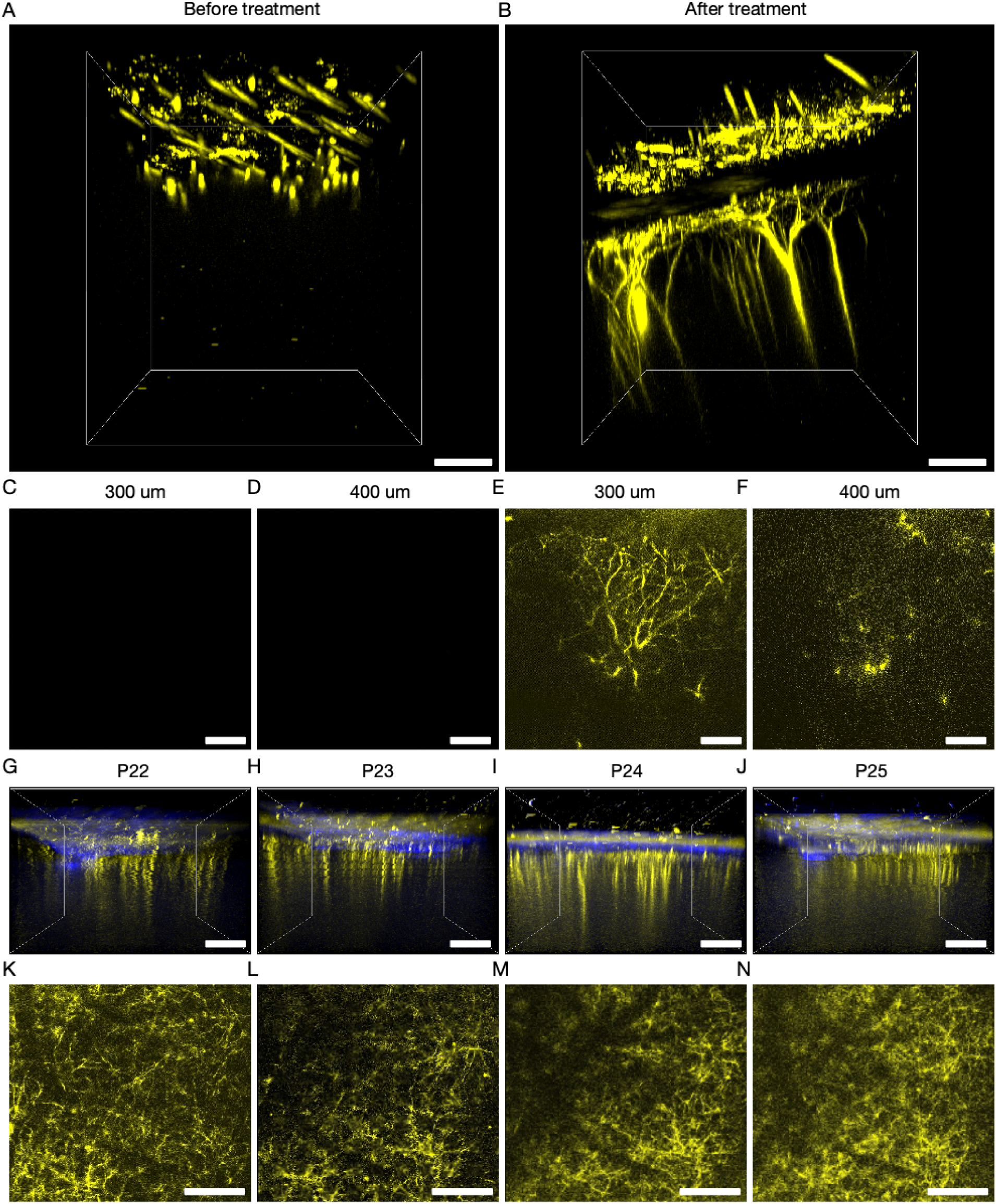
Longitudinal neuron structural imaging through transparent scalp. (*A*) 3D reconstruction of two-photon excited YFP-H fluorescence in the live mouse cortex before treatment with ampyrone. Only YFP signals in the scalp can be seen due to the scattering of the scalp. (*B*) 3D reconstruction of two-photon excited YFP-H fluorescence of the same region as *A* after achieving scalp transparency with ampyrone. (*C-D*) Images of the same brain region before treatment with ampyrone at 300 μm and 400 μm beneath the cortical surface, respectively. (*E-F*) Images of the same brain region after achieving scalp transparency with ampyrone at 300 μm and 400 μm beneath the cortical surface, respectively. (*G-J*) 3D reconstruction of two-photon excited YFP-H fluorescence of the same region in layer 1 of the primary visual area (V1) through the transparent window in the scalp on P22-P25. Yellow and blue colors indicate YFP and SHG, respectively. (*K-N*) YFP images of the same region in layer 1 of V1 at a depth of 250 μm below the surface of the scalp on P22-25. The same vascular landmarks, appearing as dark linear features, are present in all images, confirming longitudinal imaging in the same mouse brain. Scale bars are 100 μm in *A-F*, and 200 μm in *G-N*.

Cross-sectional fluorescence images at representative depths of 300 μm and 400 μm below the surface of the scalp further illustrate the contrast between the same mouse before and after achieving scalp transparency (Fig. 5*C-F*). Specifically, in the untreated condition, virtually no neuronal signals were detectable through the opaque scalp (Fig. 5*C&D*), whereas after ampyrone treatment, clear neuronal structures became visible at these depths (Fig. 5*E&F*). Notably, ampyrone induces scalp transparency without penetrating the skull. In four-week-old mice, scalp transparency alone was sufficient to enable visualization of cortical neuron structures through both the scalp and skull (see **Supporting Text**). We attribute this to the fact that, at this age, the scalp is the primary source of light scattering, while the skull remains thin enough to permit optical transmission. To the best of our knowledge, this demonstration represents the first successful imaging of YFP-expressing neurons through both the scalp and skull in juvenile mice.

The transient and reversible scalp transparency enabled repeated imaging of the same neuronal structures in the same animal for longitudinal studies. The rapid changes in brain size during early development pose challenges for conventional intravital window installations. We used ampyrone to create a transparent scalp window repeatedly, allowing cortical neuron imaging in the same Thy1-YFP-H mouse over four consecutive days, from postnatal day 22 (P22) to P25 (Fig. 5*G–N*). Single cross-sectional images taken at a fixed depth of 250 μm in layer 1 of the primary visual cortex (V1) from the scalp surface on different postnatal days revealed consistent neuronal structures in the same brain region, as evidenced by identical blood vessel features (appearing as negative contrast) across all images. This study demonstrates the feasibility of longitudinal imaging with repeatable scalp transparency *in vivo*, which could be particularly valuable for studying neurodevelopment in mice.

### Ampyrone enables functional calcium imaging through the scalp in live mice

After successfully demonstrating structural imaging of neurons in the brain through the transparent scalp, we next aimed to evaluate functional calcium imaging using GCaMP in awake mice. Specifically, we transcranially injected wild-type mice with a GCaMP8m adeno-associated virus (AAV) in the right cortical hemisphere on the day of birth (P0). Imaging was performed 21 days postnatally (P21). Similar to Fig. 5*A*, in the presence of an intact scalp, two-photon fluorescence microscopy primarily revealed autofluorescence from hair follicles, with minimal discernible cortical features underneath (Fig. 6*A*). In contrast, after rendering the scalp transparent with ampyrone, two-photon microscopy clearly resolved neuronal cell bodies in the cortex at a depth of approximately 200 μm from the surface of the scalp (Fig. 6*B*). Cross-sectional fluorescence images at a fixed depth of 170 μm from the surface of the scalp further demonstrated this contrast, with significantly improved visibility of cortical features through the transparent scalp (Fig. 6*C&D*).

**Fig. 6.**
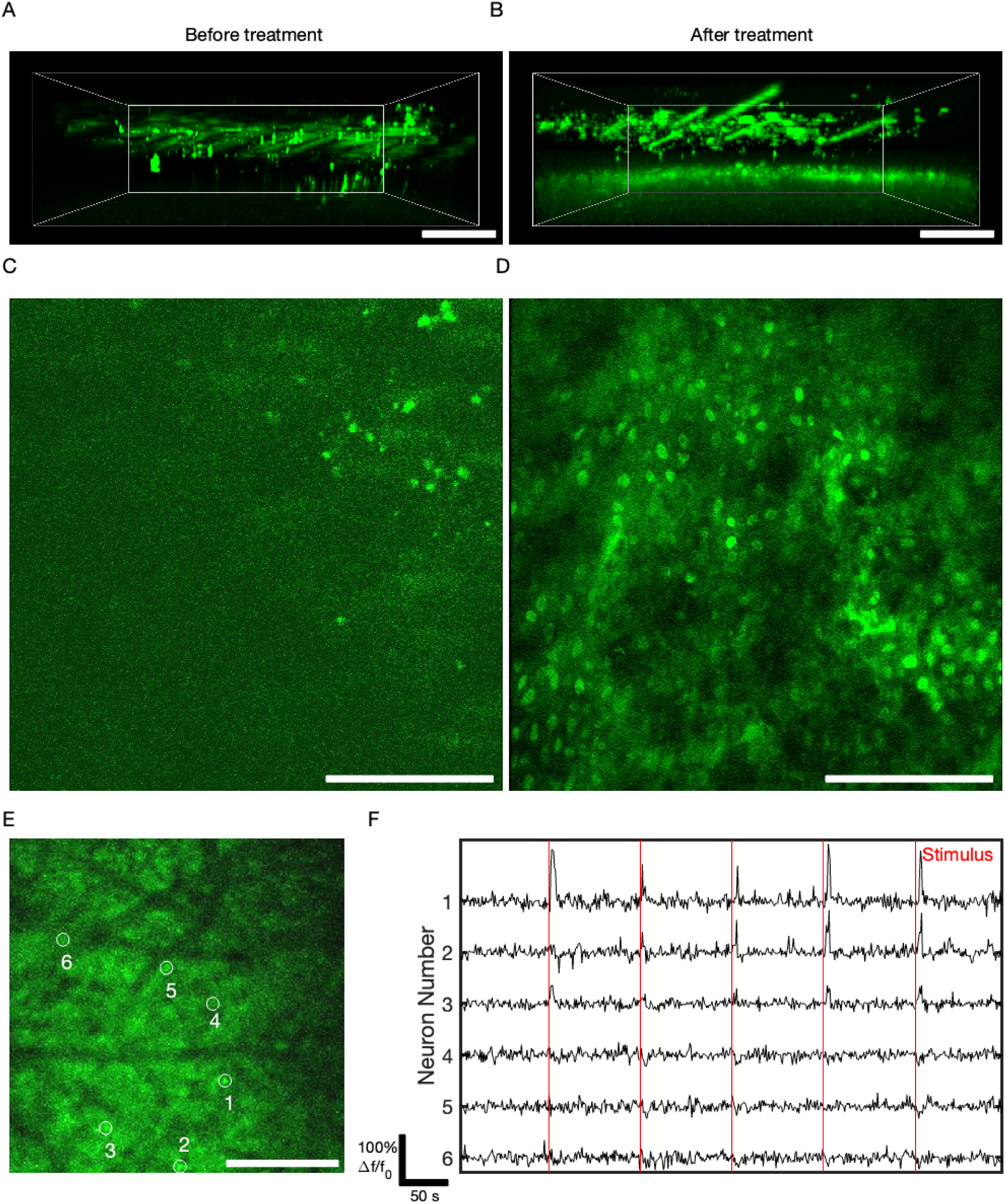
Functional neuron activity imaging with GCaMP through transparent scalp. (*A*) 3D reconstruction of two-photon excited GCaMP8 fluorescence in the mouse cortex before treatment with ampyrone. Only GCaMP8 signals in the scalp can be seen due to the scattering of the scalp. (*B*) 3D reconstruction of the same mouse cortex after achieving transparency in the scalp. (*C)* Image from the same ROI as *A* at 170 μm depth from the surface of the scalp. (*D*) Image from the same ROI as *B* at 170 μm depth showing dense cell bodies. (*E*) A representative time-frame image of the cortex of a GCaMP6f labeled mouse at 200 μm depth from the surface of the scalp. Representative neurons are labeled with white circles. (*F*) Dynamic time traces of GCaMP6f fluorescence intensity corresponding to the labeled neurons in *E*. Red lines indicate the stimulus applied to the mouse. Scale bars are 100 μm in *A-B,* 200 μm in *C-D*, and 400 μm in *E*. The different strain, larger FOV, use of awake mice, and lower laser power used for functional imaging in *E* are responsible for the lower resolution of cell bodies in *E* compared to *D*.

We next conducted time-lapse two-photon fluorescence microscopy of GCaMP6f through the transparent scalp in awake behaving head-fixed mice. Representative time-dependent traces from six selected neurons (Fig. 6*E*) showed distinct calcium transients in response to air-puff stimulation applied to the whiskers and face. Notably, three of the six neurons (neurons 1, 2, and 3) exhibited a clear stimulus response, while the remaining three (neurons 4, 5, and 6) showed minimal response (Fig. 6*F*). Importantly, the distinct responses of individual neurons ruled out motion artifacts as a source of calcium transients (Fig. S6). Fourier transform analysis of the same brain region, imaged both through the transparent scalp and after scalp removal, confirmed that imaging resolution through the transpaernt scalp was comparable to that achieved with the scalp entirely removed (Fig. S7). These experiments demonstrate that ampyrone enables functional calcium imaging of cortical neurons through the transparent scalp and skull, expanding the potential for noninvasive *in vivo* neuroimaging in awake and behaving mice.

### Proteomic analysis reveals that ampyrone does not significantly increase apoptotic markers

To further evaluate the biosafety of ampyrone-induced *in vivo* tissue transparency, we performed proteomic analysis on mice treated with ampyrone on their abdominal skin. Apoptotic markers, particularly members of the caspase and Bcl-2 family, were evaluated, as these proteins play key roles in programmed cell death. The caspase family of cysteine proteases, including Casp3, Casp6, Casp7, Casp8, and Casp9, are predominantly linked to their ability to enact or enable functions related to apoptosis (28), while Bax, a proapoptotic member of the Bcl-2 family, serves as an established indicator of apoptotic signaling (29, 30).

Both male and female mice were treated with ampyrone and allowed to recover for approximately 72 h before protein extraction from the abdominal skin, followed by sample preparation for mass spectrometry (Fig. 7*A*). Control mice underwent an identical procedure with PBS in place of ampyrone. Using a significance threshold of P < 0.05 for fold change enrichment, we observed no significant upregulation of apoptotic markers in ampyrone-treated samples compared to controls (Fig. 7*B&C*). These findings suggest that ampyrone does not induce apoptotic signaling in treated tissues, reinforcing its potential suitability for *in vivo* tissue transparency applications.

**Fig. 7.**
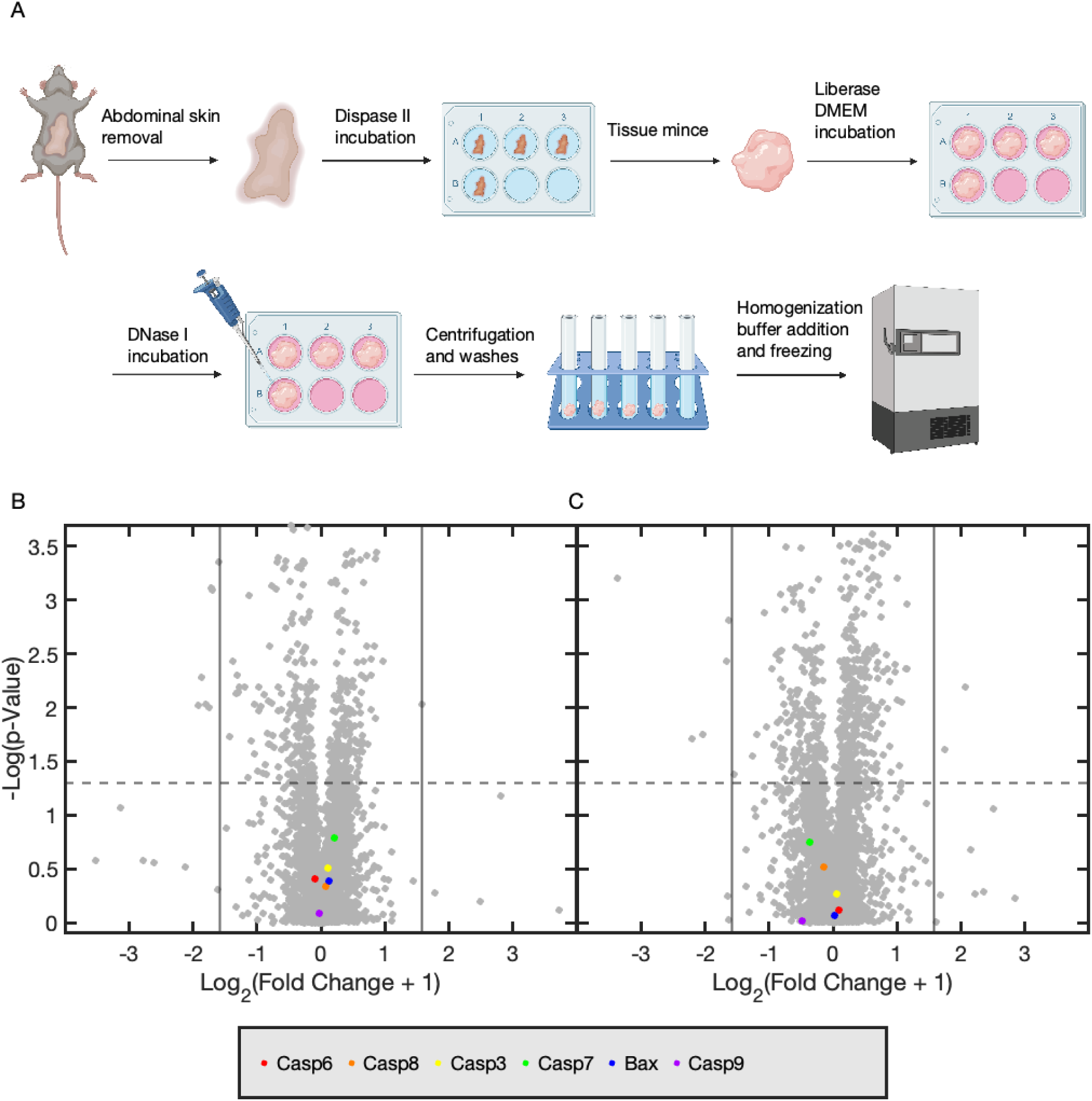
Proteomics results of mouse abdominal skin. (*A*) Schematic of protein extraction from mouse abdominal skin following a 72 h recovery period post treatment with ampyrone. (*B*&*C*) Volcano plot depicting the female (*B*) and male (*C*) mouse proteomics results with potential apoptotic markers highlighted. Dashed, horizontal lines correspond to the 0.05 P-value threshold for significance and vertical lines correspond to fold changes of ±2, which corresponds to a Log(Fold Change) of Log_2_(2+1). The fold change is defined as (final value – original value)/(original value).

## Discussion

In this study, we have demonstrated that *in vivo* optical tissue transparency, previously limited to red visible wavelengths in our earlier work (14), can now be extended across the entire visible spectrum. This advancement significantly broadens the applicability of the *in vivo* transparency approach, making it compatible with a wide range of fluorescent proteins and other fluorescent probes throughout the visible spectrum. Leveraging this development, we successfully performed both longitudinal and functional imaging of neurons in the mouse cortex using YFP and GCaMP, respectively, through the intact scalp and skull of live mice.

While previous studies have achieved two-photon fluorescence microscopy through the skull (31) and even more advanced three-photon transcranial imaging (32), to our knowledge, this study is the first demonstration of cellular-resolution imaging of neuronal structures and calcium activity through both the scalp and skull. Although our current work has focused on yellow and green fluorescent proteins, red fluorescent proteins such as mCherry, tdTomato, and RCaMP should also be fully compatible, enabling multiplexed brain imaging with multiple fluorophores in distinct color channels through ampyrone-treated transparent tissue.

Beyond expanding optical transparency to the full visible spectrum in living tissue, we have also demonstrated the reversibility of our transparency approach in the mouse abdominal skin and shown that scalp transparency can be achieved while the mouse remains awake and behaving (Movie S1). This reversible and transient nature of ampyrone treatment ensures biosafety, as supported by blood chemistry, hematology (Fig. S4), skin histology (Fig. S5), and proteomics analyses (Fig. 7). Additionally, it enables chronic imaging by allowing the same skin region to be rendered transparent repeatedly over multiple days (Fig. 5).

Although our study indicates excellent recovery following ampyrone treatment, long-term effects still need to be thoroughly investigated in mice before progressing to larger animal models. Additionally, we aim to extend our approach beyond skin transparency to other internal organs, with a particular focus on directly rendering the brain transparent in future research. However, achieving optical transparency in internal organs presents greater challenges due to the necessity of more invasive delivery methods and the increased vascularization, which may lead to rapid clearance of RI-matching agents.

Future advancements in absorbing molecule design could further enhance this approach. Fluorophores with stronger ultraviolet resonances and broader absorption spectra could improve transparency efficiency, allowing for even lower concentrations to achieve effective RI matching in tissue. Alternatively, engineered dyes with narrow absorption profiles could be optimized for operation near specific resonances, enhancing the precision and efficacy of optical tissue transparency.

## Materials and Methods

### Chemicals and materials

Ampyrone was purchased from Sigma-Aldrich (33528-100G-R, batch BCCK4073, Sigma-Aldrich, Burlington, Massachusetts, USA) and stored in an MBRAUN Unilab nitrogen purged glove box fridge (14-091, MBRAUN, Garching, Germany) at −20°C before use. Powder aliquots were stored under vacuum at room temperature for <1 week before immediate use and were covered in foil to prevent exposure to ambient light.

For *ex vivo* experiments, ampyrone solutions were prepared by dissolving 200 mg, 400 mg, 600 mg, or 800 mg of ampyrone in 1 mL of deionized water at 40°C for 10 min, resulting in concentrations of 17% w/w, 29% w/w, 38% w/w, and 44% w/w, respectively. Occasional shaking was used to ensure complete dissolution. After preparation, ampyrone solutions were stored at ambient conditions and used within 12 h; any unused solution was discarded to maintain experimental consistency.

For *in vivo* experiments, ampyrone solutions were prepared by dissolving 600 mg of phenazone in 1 mL of deionized water at 40°C for 10 min to achieve a 38% w/w concentration. Occasional shaking was used to ensure complete dissolution. The solutions were stored at ambient conditions and used within 12 h; any remaining solution was discarded to maintain experimental consistency.

Phenazone (A5882-100G, Sigma-Aldrich, Burlington, Massachusetts, USA), glycerol (G5516-1L, Sigma-Aldrich, Burlington, Massachusetts, USA), and dextran (MW: 40 kDa, 31389-25G, Sigma-Aldrich, Burlington, Massachusetts, USA) were all purchased from Sigma-Aldrich. Sucrose (S5-500, Fisher Scientific Company, Pittsburgh, Pennsylvania, USA) was purchased from Fisher Scientific Company. Deionized water was obtained from 18.2 MOhm deionized Milli-Q Integral 10 (7003, EMD Millipore, Burlington, Massachusetts, USA) and used as the solvent for all experiments.

### Spectroscopic ellipsometry

All spectroscopic ellipsometry experiments were performed on a Horiba Jobin Yvon UVISEL (2015 model, Horiba Jobin Yvon IBH Limited, Irvine, California, USA) spectroscopic ellipsometer. The bulk complex refractive index was measured for each solution with a volume of 2.5 mL. Tissue-Tek Cryomolds (4557, Sakura Finetek, 25 mm x 20 mm x 5 mm, Torrance, California, USA) were used to hold the solution, secured to the UVISEL stage with a built-in vacuum chuck. To eliminate reflections from surfaces other than the top liquid surface of the sample, 320-grit sandpaper (SiC A-99, Partsmaster 881-5-0320, Dallas, Texas, USA) was attached to the bottom of the cryomold using double-sided tape (3M 34-8724-5691-7, Saint Paul, Minnesota, USA). This ensured that only the bulk complex refractive index of the liquid sample was measured, without the need for multilayer postprocessing of the raw ellipsometric data. Spectra were taken from 250 nm to 850 nm with a 2 nm step size and a 200 ms dwell time.

### UV-Visible spectrophotometry

All absorption and transmission experiments were performed on a Thermo-Scientific Evolution 350 UV-Visible spectrophotometer (Thermo-Scientific, Waltham, Massachusetts, USA). Fused quartz cuvettes with 1 mm pathlength (Aireka Scientific Co Ltd, Hong Kong, China) were used for all liquid measurements. The UV-Visible transmission spectra of ampyrone, phenazone, and tartrazine were measured at concentrations of 100 mg/mL for each compound in Fig. 1A. For transmission measurements of mouse skin samples, the samples were placed on glass slides (3” x 1” x 1 mm, 12-544-1, Fisher Scientific, Pittsburgh, PA, USA) and secured with double-sided tape (34-8724-5691-7, 3M, Saint Paul, MN, USA) to built-in cuvette holders (Standard Cell Holder for Thermo 350, Thermo-Scientific, Waltham, MA, USA) within the UV-Vis spectrophotometer. The double-sided tape was placed around the optical window of the cuvette holder to avoid interfering with the spectrometry measurements. Max Bond Super Glue (Krazy Glue, High Point, North Carolina, USA) was used to glue the corners of the skin if the tissue failed to adhere. Spectra were taken from 190 nm to 1100 nm with 0.25 s dwell time, 1 nm step size, and 4 nm bandwidth.

### Scattering phantom imaging

For phantom images in Fig. 2*A*, scattering phantoms measuring 15 mm × 15 mm × 5 mm were prepared by adding 0, 400, 800, 1200, or 1600 mg of ampyrone (corresponding to % w/w concentrations of 0%, 17%, 29%, 38%, and 44%, respectively) along with 20 mg of SeaPlaque agarose (50101, Lonza, Rockland, MO, USA) into a 20 mL scintillation vial (DWK986546, Sigma-Aldrich, Burlington, MA, USA). 2 mL of 10 mg/mL 1 μm silica particle suspension (SISN1000, nanoComposix, San Diego, California, USA) was then added to each vial. The suspension was heated at 80°C for 10 min and then thoroughly mixed using a vortex. It was then heated for another 10 min at 80°C and mixed again with a vortex. Next, 1.2 mL of each suspension was cast into Tissue-Tek Cryomolds (15 mm × 15 mm × 5 mm, 4557, Sakura Finetek, Torrance, CA, USA) and sealed with a coverslip to ensure a phantom thickness of exactly 5 mm. The samples were chilled on ice for 10 min to set the gel before imaging. The samples were placed on a 300 lumen LED light board (A4 LED Light Board, Comzler, Dongguan, Guangdong, China) with an attached 1 x 1 mm grid pattern printed on a transparency film (Gwybkq Transparency Film, 8.5 x 11 Inches, Amazon, Seattle, WA, USA). Imaging was done with a Canon EOS Rebel T6 Digital SLR camera (DS126621, Canon, Tokyo, Japan) equipped with a Canon EFS 18-55 mm MACRO 0.25 m/0.8 ft lens (226064565741, Canon, Tokyo, Japan).

### USAF resolution target imaging

For USAF resolution target imaging, circularly shaped scattering phantoms were used. Specifically, scattering phantoms with a 2 mm thickness and a 6 mm diameter were prepared by stacking two CultureWell reusable gaskets (CW-8R-1.0, Grace Bio-Labs, Bend, Oregon, USA) onto glass slides (Fisherbrand premium plain glass microscope slides, 3 in x 1 in x 1 mm, 12-544-1, Fisher Scientific, Pittsburgh, Pennsylvania, USA). Scattering phantoms were prepared by adding 0, 20, 40, 60, or 80 mg of ampyrone (corresponding to concentrations in % w/w of 0%, 17%, 29%, 38%, and 44%, respectively) with 1 mg of SeaPlaque agarose (50101, Lonza, Rockland, Missouri, USA) to a 1.8 mL scintillation vial (03338AA, Fisher Scientific, Pittsburgh, PA, USA). 100 μL of 10 mg/mL 1 μm silica particle suspension (SISN1000, nanoComposix, San Diego, California, USA) was then added to each vial. The suspension was heated at 80°C for 10 min and then thoroughly mixed using a vortex. It was then heated for another 10 min at 80°C and mixed again with a vortex. The suspension was then cast into the silicon gaskets. Each gasket can hold 8 separate wells, with each well containing a different sample with a different ampyrone concentration. Another glass slide was placed on top to seal the samples in each of the gaskets. The samples in the gaskets were chilled on ice for 10 min to set the gel before imaging.

A negative 1951 USAF resolution test target (R3L3S1N, Throlabs, Newton, New Jersey, USA) was secured to the stage of a Leica DM2700M microscope (Leica, Wetzler, Germany) with a 10X (0.25 BD N PLAN EPI) objective (Leica, Wetzler, Germany). The USAF target and scattering phantom sample being imaged were centered in the field of view of the microscope. The target was secured with tape to prevent movement while the gaskets were moved to center different wells with different phantom samples. Images were acquired with a Hamamatsu C11440-22CU digital camera (Hamamatsu, Shizuoka, Japan). For Fig. 2*C*, these images were digitally cropped such that only the group 6-7 elements (the two groups with the smallest and second smallest features) of the USAF target were visible. The following filters were used: 450/10 (FBH450-10, Thorlabs, Newton, New Jersey, USA), 525/45 (BrightLine Fluorescence Filter 525/45, Semrock, Northbrook, Illinois, USA), 600/10 (FB600-10, Thorlabs, Newton, New Jersey, USA), 680/10 (FB680-10, Thorlabs, Newton, New Jersey, USA), and 785LP (Edgebasic Long Wave Pass 785, Semrock, Northbrook, Illinois, USA). To quantify the contrast of the USAF target through the scattering phantoms, the modulation transfer function (MTF) was calculated as described below in the “Modulation transfer function (MTF) calculation” section.

### Modulation transfer function (MTF) calculation

The MTF is one common metric for quantifying the contrast of an optical system at specific spatial frequencies. It quantifies the ability of a system to capture detailed structures in an image and is based on the spatial frequency of periodic patterns such as line or sine waves (33–35). These features are measured in units of line pairs per millimeter (lp mm^−1^). The 1951 USAF test target is one of the most popular image targets for quantifying the MTF. The MTF itself can be calculated by first extracting the normalized difference between the maximum and minimum transmitted light intensities at a specific spatial frequency from an image of a USAF test target. The MTF can then be calculated as:

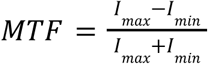

Where *I_max_*is the maximum intensity of a specific spatial frequency feature in the USAF target and *I_min_* is the minimum intensity of the same feature.

### Vertebrate animal subjects

Mice were group-housed on a 12 h:12 h light:dark cycle in the Stanford University Veterinary Service Center (VSC) with food and water provided *ad libitum*. All animal experiments conducted were approved by Stanford University’s Administrative Panel on Laboratory Animal Care (APLAC) in accordance with Public Health Service Policy on Humane Care of Laboratory Animals guidelines and were approved by the Institutional Animal Care and Use Committees of Stanford University.

### Achieving optical transparency in *ex vivo* mouse skin

C57BL/6J mice (male and female, 8-10 weeks old, 20-30 g) from Jackson Labs (Sacramento, California, USA) were used for demonstrating optical transparency in *ex vivo* mouse skin tissue. The mice were euthanized using carbon dioxide via inhalation, in accordance with the APLAC protocol, followed by cervical dislocation to ensure death. The abdominal skin was then surgically removed after depilation by incising an approximately 10 mm x 20 mm rectangle of abdominal skin and peritoneum from the center of the abdomen. The peritoneum was subsequently removed from the skin. One mouse was used for a single skin sample. Skin samples were glued by the corners onto glass coverslips with Max Bond Super Glue (Krazy Glue, High Point, North Carolina, USA) and placed in a 12 well plate, with one sample per well. The 12 well plate was placed on a 300 lumen LED light board (A4 LED Light Board, Comzler, Dongguan, Guangdong, China) with an attached 1 x 1 mm grid pattern printed on transparency film (Gwybkq Transparency Film, 8.5 x 11 Inches, Amazon, Seattle, WA, USA) in an orbital shaker (MaxQ 4000, Thermo Scientific, Waltham, Massachusetts, USA) set to 40°C. Samples were soaked in aqueous solutions of ampyrone at concentrations of 17%, 29%, 38%, or 44% w/w for 24 h at 40°C without shaking. Images before and after soaking were taken with a digital camera (DCU223M, ThorLabs, Newton, New Jersey, USA) using a Xenon variable aperture f/0.95 25 mm focal length lens (Model 13238999, Schneider-Kreuznach, Bad Kreuznach, Germany) at 4 frames per second (FPS), 200 ms exposure, and f/11 aperture. For obtaining the T% value for each sample, a blank coverslip identical to the ones used to hold the abdominal skin was used as the baseline (i.e., T% = 100%) before each measurement. The blank coverslip was attached to the cuvette holder as described above. For obtaining the T/T_original_ value for each sample, the T% value for the sample after soaking was divided by the T% value for the same sample before soaking.

### Achieving optical transparency for *in vivo* abdominal imaging

C57BL/6J and BALB/cJ mice (male and female, 3-4 weeks old, 9-15 g) from Jackson Labs (Sacramento, California, USA) were used for *in vivo* abdominal imaging. The mice were anesthetized (Animal Anesthesia Vaporizer, RWD, Sugar Land, Texas, USA) with Isospire^TM^ isoflurane (Dechra Veterinary Products, Overland Park, Kansas, USA) and secured to a heating pad (731-500-000R, Sunbeam, Boca Raton, Florida, USA). Mice were subcutaneously dorsally injected with 20 μL saline per gram of body weight (0.9% sodium chloride injection, USP, Hospira Inc, Lake Forest, Illinois, USA) to maintain hydration. Abdominal fur was removed with Nair hair removal body cream (NRSL-22339-07, Church and Dwight Co., Ewing, New Jersey, USA). Specifically, cotton tipped applicators (803-WC Hospital, Puritan Medical Products Company LLC, Guilford, Maine, USA) were used to first apply a generous amount of Nair to the fur. The Nair-covered fur was massaged lightly with the cotton tipped applicators for 5 min. The cotton tipped applicators were then used to remove the fur with wiping motion. Sterile alcohol prep pads (Cat. No. 22-363-750, FisherBrand, Pittsburgh, Pennsylvania, USA) were then used to remove the remaining fur and Nair through wiping. The abdominal skin was then exfoliated with Nair for another 5 min with gentle massaging. After removing Nair with alcohol wipes, approximately 0.1 mL of the ampyrone solution (38% w/w, aqueous) was immediately applied with a cotton-tipped applicator before the residual alcohol on the skin had fully dried. The solution was then lightly massaged over an approximately 1 cm^2^ area of skin for 5 min. It is estimated that only 20% or less of the ampyrone solution will make it into the skin, with the rest falling off or being absorbed into the cotton tipped applicator. The mouse belly was gently pressed with a glass slide (FisherBrand plain microscope slides, 75 x 50 x 1.0 mm, Pittsburgh, Pennsylvania, USA) to flatten features for imaging. Imaging was done with a Canon EOS Rebel T6 Digital SLR camera (DS126621, Canon, Tokyo, Japan) equipped with a Canon EFS 18-55 mm MACRO 0.25 m/0.8 ft lens (Canon, Tokyo, Japan) illuminated by a portable photography light (MOLUS G60 60W portable photography light, Zhiyun, Quangxi, China) set to 6 W illumination power (10%) and 5500 K color temperature. Both the camera and illumination were linearly polarized and set at angle with respect to each other to eliminate reflection from the gel or glass slide.

After imaging, the mouse abdomen was rinsed with warm 5 mL PBS (PBS, 1X, Corning Inc., Corning, New York, USA) gently with a transfer pipette (Cat. No. 13-711-9AMMD, FisherBrand, Pittsburgh, Pennsylvania, USA) to extract the skin-absorbed ampyrone and reverse the transparency effect. Delicate task wipers (Kimberly-Clark, Irving, Texas, USA) were used to prevent the fur from getting wet to prevent hypothermia or infection. We estimate that ∼80% of the skin-absorbed ampyrone was removed from the skin during this step. The mouse abdomen was then wiped with alcohol prep pads (Cat. No. 22-363-750, FisherBrand, Pittsburgh, Pennsylvania, USA) one more time before eye lubricant gel (Cat. No. B000URVDQ8, Genteal, Fort Worth, Texas, USA) was applied to hydrate the skin. A tough bioadhesive hydrogel (Sonopatch AD48, Sonologi, Palo Alto, California, USA) with a minimum 1 cm x 1 cm area was cut and placed onto the abdomen with the adhesive side touching the skin. The hydrogel pad was placed on the skin for 1 min with gentle pressure to ensure adhesion. The mouse abdomen was then wrapped with surgical tape (B014SPIK22, 3M Durapore 1” x 30’, Saint Paul, Minnesota, USA) to protect the hydrogel. It is essential that the tape be wrapped around itself such that the adhesive side of the tape touches itself. If the tape does not have two adhesive sides touching, it will come off when the mouse becomes active. The mouse was kept in a cage with bedding set on top of a heating pad (731-500-000R, Sunbeam, Boca Raton, Florida, USA), but with only half of the cage on the heating pad so that the mouse had the option to stay on the room temperature side of the cage.

For longitudinal abdominal skin imaging, the same recovery procedure as above was repeated for every day of imaging. The surgical tape and hydrogel were removed immediately before imaging, followed by reapplication of eye gel, hydrogel, and surgical tape. If imaging was not required every day, the hydrogel and surgical tape were only replaced after the mouse had damaged the surgical tape.

### Achieving optical transparency for two photon excitation fluorescence imaging in the live mouse brain

Thy1-YFP-H mice (male and female, 3-4 weeks old, 9-15 g, C57BL/6J [strain #000664] x B6.Cg-Tg(Thy1-YFP)HJrs/J [strain #003782]) from Jackson Labs were used for structural imaging of the brain. Mice were anesthetized with isoflurane with a custom 3D-printed anesthetic mask (*SI Appendix,* Dataset S1) and secured with a head holder (SGM-4, Narishige, Amityville, New York, USA). Mouse body temperature was maintained with a silicone rubber fiberglass flexible heater (SRFRA-8/*, Omega Engineering, Norwalk, Connecticut, USA). Mice were subcutaneously injected dorsally with 20 μL saline per gram of body weight (0.9% sodium chloride injection, USP, Hospira Inc, Lake Forest, Illinois, USA) to maintain hydration. The scalp of the mouse was depilated and then exfoliated with Nair following the same procedure as above. Following exfoliation, ampyrone was applied to the scalp similarly to the abdomen until the scalp becomes visibly transparent to reveal the features in the skull such as the lambda, bregma, and other bone plate borders. A coverslip was secured to the top of the scalp above the visual cortex with a custom 3D-printed coverslip holder (*SI Appendix,* Dataset S2) to isolate the objective from the remaining ampyrone solution, which acted as an index-matching medium between the coverslip and the transparent scalp. The coverslip holder provides pressure to the coverslip without obscuring the optical path of the microscope. Two-photon excitation fluorescence imaging was done on a Prairie Ultima IV Two-photon *In Vivo* Microscope (Bruker, Billerica, Massachusetts, USA). A water immersion objective (20X Olympus XLUM Plan Fl W, 0.95 NA and 2.0 mm working distance, Olympus, Tokyo, Japan) was used. 920 nm excitation was provided by a Mai Tai HP Deep See Ti-sapphire laser (Spectra Physics, Milpitas, California, USA) while emission was collected through a 525/50 nm bandpass filter. Pixel dwell time was 1.2 μs. Image size for Z-stacks was 512 x 512 pixels per frame. Field of view (FOV) for each image in the Z-stacks was 598 μm x 598 μm. Manual gain compensation was used for all Z-stacks.

For longitudinal YFP imaging, the same procedure was done as above except for modified imaging conditions and the recovery procedure (See *SI Appendix* supporting information text for details). Specifically, mice were imaged while awake with the heating pad replaced with a treadmill to enable free mouse movement during imaging. To minimize motion artifacts and facilitate the consistent identification of the same blood vessel features across different days, a 10X air objective (10X Olympus UPlanFL N, 0.30 NA and 10 mm working distance, Olympus, Tokyo, Japan) was used. The stereotactic coordinates for the center of the FOV (in anterior-posterior (AP), medial-lateral (ML), dorsal-ventral (DV) coordinates with bregma set as zero) are (−3.2 mm, −2.3 mm, 0.2 mm). This corresponds to layer 1 of the primary visual cortex (V1). In addition to collecting the YFP emission with a 525/50 bandpass filter, the second harmonic generation (SHG) signal from collagen primarily in the scalp and skull was collected with a 460/50 bandpass filter to enable better z reference. Pixel dwell time was 1.2 μs and image size for Z-stacks was 512 x 512 pixels, with a FOV of 1.170 mm x 1.170 mm. Manual gain compensation was used for both the YFP and SHG channels independently.

Two mouse lines were used for calcium imaging. The first line was male and female Fostm2.1(icre/ERT2)Luo/J mice (Jax strain #: 030323) injected with 0.5-1 μL GCaMP8 AAV (AAV1-syn-jGCaMP8m-WPRE, titer: 2.4×10^13^ vg/mL, Addgene # 162375-AAV1) into the center of the right hemisphere at P0 (0 days after birth). This line was used in Fig. 6*A-D*. The second line was male and female GCaMP6f mice (B6J.Cg-Gt(ROSA)26Sortm95.1(CAG-GCaMP6f)Hze/MwarJ [strain #028865] x B6.Cg-Tg(Camk2a-cre)T29-1Stl/J [strain #005359]). This line was used in Fig. 6*E*. The Fostm2.1(icre/ERT2)Luo/J mice were 22 days old (P22) and the GCaMP6f mice were between 16 and 23 days old (P16 and P23). All mice were within the weight range of 8-15 g. The 20X objective and the same imaging parameters used for non-longitudinal YFP imaging were applied in Fig. 6*A*-*D* in anesthetized mice. For Fig. 6*E*, different imaging parameters were used to capture the time-series data in awake mice. Specifically, to obtain the best firing activity and stimulus response, mice for time-series GCaMP imaging were imaged while awake, using the same awake imaging setup mentioned above for the longitudinal YFP imaging. The 10X objective described above was used to minimize motion artifacts and ensure at least some active and visible neurons were in the FOV. The image size was 256 x 256 pixels, dwell time was 12.4 μs, recording frame rate was 0.987 FPS, and FOV was 1.170 mm x 1.170 mm. The approximate stereotactic coordinates of the center of the FOV in Fig. 6*E* (in AP, ML, DV coordinates with bregma set as zero) are (−1.2 mm, −0.9 mm, 0.3 mm). This location corresponds to layer 1 of the retrosplenial cortex (RSC). Vascular landmarks were checked with vascular maps of the mouse brain in literature to verify location (36–38). To provide stimulus, a hose was attached to the imaging setup and air was manually generated using a high pressure dust remover (Decon Labs Inc., Prussia, Pennsylvania, USA) to provide air pressure to both the mouse whiskers and face, promoting a running response on the treadmill. The RSC is responsible for integrating sensory and nonsensory information, with correlations between running speed and activity in the RSC reported (39). Full setups for both longitudinal and functional imaging are shown in Fig. S8. For both YFP and GCaMP imaging, the mice were less than 28 days old (P28) to ensure that the skull was not too thick for through-skull imaging. The reasoning behind this choice is that, although ampyrone effectively eliminates scattering caused by the scalp, it cannot penetrate or render the skull transparent.

For 3D reconstruction from the z-stacks, IMARIS was used, with the 3D reconstructions processed by applying background correction, gamma correction, and a gaussian filter, followed by false coloring as well as the adjustment of brightness and contrast. To process the calcium data, images were aligned using the Enhanced Correlation Coefficient (ECC) algorithm from OpenCV (40). A translation motion model was used to align images with respect to the first image. A 50-element sliding median was used to calculate the baseline for baseline correction. To quantify time series changes in fluorescence intensity, we defined *f* as the time-dependent baseline corrected calcium signal and *f*_0_ as the average signal from a low neural activity time window of the baseline-corrected signal when no stimulus was applied. Then Δ*f* was calculated as Δ*f* = *f* − *f*_0_. The motion-corrected video used for Fig. 6*F* is attached in *SI Appendix,* Movie S2.

### Protein extraction from mouse skin

Age-matched C57BL/6J mice (16 weeks old, 20-30 g; five males and females per group) were used for protein extraction experiments. Mice were massaged on the abdomen with the 38% w/w ampyrone aqueous solution for 5 min, while the control group received the same treatment with phosphate buffered saline (PBS, 1X, Corning Inc., Corning, New York, USA). Following the treatment, all mice were subjected to bioadhesive hydrogel application under the same conditions as abdominal imaging described above in “Achieving optical transparency for *in vivo* abdominal imaging**”**. Following a 72 h recovery period, all mice were humanely euthanized by cervical dislocation and the abdominal skin was excised from the mice, ensuring that the abdominal wall was not attached. Prior to the digestion (41), the excised skin was washed twice with 1 mL of 1X PBS, followed by dermal-side-down soaking in Dispase II solution (Dispase II, Millpore Sigma D4693) at 5 mg/mL for 30-40 min at 37°C. The dermis was finely chopped and spread across a Petri dish, then subjected to enzymatic digestion in Liberase™ (Liberase™ Research Grade, Millipore Sigma #5401119001) at 1 mg/mL diluted in DMEM (DMEM, high glucose, pyruvate, Gibco, #11995073) for 20-30 min at 37°C with gentle agitation. DNase I (Deoxyribonuclease I from bovine pancreas, Millipore Sigma DN25) at 1 mg/mL was added to the mixture and incubated for 15 min with agitation ceasing after the DNase I was added. With DNase I incubation, the digested tissue was centrifuged at 4000 relative centrifugal force (RCF) for 5 min, supernatant was removed, and the pellet was washed twice with 1 mL of 1X PBS before resuspension in 500 mL of homogenizer buffer (T-PER Tissue Extraction Reagent, Thermo Scientific #78510) containing a 1:100 protease inhibitor cocktail (Protease Inhibitor Cocktail, Millipore Sigma P8340). Samples were frozen at −80°C for 12 h, then centrifuged at at 5000 RCF for 10 min and the supernatant was transferred into low-bind protein tubes (Protein LoBind 1.5 mL Polypropylene Snap Cap Microcentrifuge Tubes, Eppendorf, #13-698-794). Protein concentration was measured by bicinchoninic acid (BCA) protein assay (Pierce BCA Assay, Thermo Scientific #23225) and the samples were prepared for mass spectrometry.

### Tandem mass spectrometry sample preparation

20 μg resuspended epidermis protein extract for each of the samples were reduced in a solution of 5% v/v sodium dodecyl sulfate (15553027, Thermo Fisher Scientific, Waltham, Massachusetts, USA) with 10 mM dithiothreitol (Bio-Rad, 1610611, Hercules, California, USA) and incubated at 95°C for 10 min. Samples were then allowed to cool to room temperature and then alkylated using 40 mM 2-chloroacetamide (C0267-500G, Sigma-Aldrich, St. Louis, Missouri, USA) and incubated for 1 h at room temperature. The alkylation reaction was quenched by adding 85% m/v phosphoric acid (W290017-1KG-K, Millipore Sigma, Burlington, Massachusetts, USA) to a final concentration of 1.2% v/v phosphoric acid in the alkylated protein solution. 700 μL (7X volume) of S-trap bind wash buffer, 100 mM triethylammonium bicarbonate buffer (TEAB, 60044974, Fisher Scientific, Hampton, New Hampshire, USA), and 90% v/v methanol (A452-4, Fisher Scientific, Hampton, New Hampshire, USA), was then added to the alkylated sample. Samples were then loaded onto a micro-S-trap-column (C02-micro-80, Protifi, Fairport, New York, USA) in 175 μL increments and spun at 4000 RCF for 20 s for each loading cycle. Sample caught on the column was then washed three times using the S-trap bind wash buffer with spins of 4000 RCF for 30 s to pass the buffer over the column. Mass spectrometry-grade trypsin (Promega V5113, Madison, Wisconsin, USA) was then diluted in 50 mM TEAB buffer (60044974, Fisher Scientific, Hampton, New Hampshire, USA) to a concentration of 0.04 mg/mL. 25 μL of the diluted trypsin solution, or 1 μg of trypsin, was loaded onto each column to make a final trypsin:protein ratio of 1:20. Samples were then incubated at 47°C for 90 min. The column was then washed using 50 mM TEAB followed by 0.2% v/v formic acid (28905, Thermo Fisher Scientific, Waltham, Massachusetts, USA) in water with a spin at 1000 RCF for 60 s between each wash. Digested samples were then eluted by adding 40 μL of a solution of 50% v/v acetonitrile (A955-212, Fisher Scientific, Hampton, New Hampshire, USA), 0.2% v/v formic acid, and 50% v/v water then spun at 4000 RCF for 60 s. Samples were then dried using vacuum centrifugation (CentriVap Complete Vacuum Concentrator, Labconco #7315023, Kansas City, Missouri, USA) for 16 h. The samples were resuspended in 0.2% (v/v) formic acid diluted in water immediately prior to mass spectrometry data acquisition.

### Tandem mass spectrometry data acquisition

All samples were resuspended in a 0.2% v/v formic acid diluted in water at 1 μg/μl, and 1 μL (1 μg) of each sample was added to a new tube to make a pooled sample (42) for the six gas phase fractionation injections. 1 μL (1 μg) of the total peptide solution was injected in the column for each sample. Peptides were separated over a 25 cm EasySpray reversed phase LC column (75 μm inner diameter packed with 2 μm, 100 Å, PepMap C18 particles, Thermo Fisher Scientific, Waltham, Massachusetts, USA). The mobile phases (A: water with 0.2% v/v formic acid and B: acetonitrile with 0.2% v/v formic acid) were driven and controlled by a Dionex Ultimate 3000 RPLC nano system (ULTIM3000RSLCNANO, Thermo Fisher Scientific, Waltham, Massachusetts, USA). Gradient elution was performed at 300 nL/min. Mobile phase B was increased from 1 to 5% v/v over 6 min, followed by a gradual increase to 25% v/v by 60 min, and ending with a ramp to 90% v/v B at 71 min, and a wash at 90% v/v B for 5 min. Flow was then ramped back to 1% v/v B over the course of 1 min, and the column was re-equilibrated at 1% v/v B for 15 min, for a total analysis of 90 min. Eluted peptides were analyzed on an Orbitrap Fusion Tribrid MS system (Thermo Fisher Scientific, Waltham, Massachusetts, USA). Precursors were ionized using an EASY-Spray ionization source (Thermo Fisher Scientific, Waltham, Massachusetts) source held at +2.2 kV compared to ground, and the column was held at 40 °C. The inlet capillary temperature was held at 275 °C. For gas phase fractionation injections, a pooled sample was injected six times and acquired using staggered-window injections with m/z windows of 395 to 505 m/z, 495 to 605 m/z, 595 to 705 m/z, 695 to 805 m/z, 795 to 905 m/z, and 895 to 1005 m/z. For all staggered-window injections a 4 m/z precursor isolation window was used. MS/MS scans were collected using high-energy collisional dissociation at 33 normalized collision energy and mass analysis was performed in the Orbitrap using a resolution of 30,000 while scanning from 120–2000 m/z. Individual samples were collected using single-injection data independent acquisitions (DIA) with survey scans of peptide precursors collected in the Orbitrap from 385-1015 m/z with an automatic gain control target of 400,000, a maximum injection time of 55 ms and a resolution of 60,000. An isolation window of 16 m/z was used to select precursor ions with the quadrupole. MS/MS scans were collected using high-energy collisional dissociation at 33 normalized collision energy and mass analysis was performed in the Orbitrap using a resolution of 30,000 while scanning from 120–2000 m/z.

### Mass spectrometry database search and quantitation

Proteins were identified with DIA-NN (43) and MSFragger 4.0 (44) software on the Fragpipe GUI version 21.0 (45), searching against the UniProt (46) mouse reviewed proteome in addition to common contaminants. Methionine oxidation (+15.995 Da) and N-terminal acetylation (+42.011 Da) were included as variable modifications, while carbamidomethylation of cysteine (+57.021 Da) was included as a fixed modification. Precursor ion search tolerance was set to 20 ppm and fragment ion mass tolerance was set to 20 ppm. Peptide and protein identifications were thresholded at a 1% false discovery rate using a target-decoy method (47). Proteins were quantified and normalized by label-free quantitation using the DIA-NN software. For quantitative comparisons, protein intensity values were log transformed, and missing values for proteins were imputed from a normal distribution with a width of 0.3 standard deviations and a down shift value of 1.8 standard deviations in Perseus version 2.0 (48). A two-sided t-test was additionally done using Perseus with false discovery rate (FDR) correction performed using permutation-based FDR with 250 randomizations. Fold change was calculated in comparison with a PBS-treated control and log2 transformed for the volcano plot.

### Animal subject preparation for tissue collection and histopathology

Age-matched C57BL/6J (male and female, 16 weeks, 20-30 g) mice were used for all experiments. 8 mice (4 male and 4 female mice) were depilated, exfoliated, treated with ampyrone in the abdominal skin, and subsequently reversed for the transparency effect as described in the “Achieving optical transparency for *in vivo* abdominal imaging” section. The mice were monitored and given the same recovery treatment with hydrogel as described above. The mice were then submitted to the Stanford Veterinary Service Center Necropsy Lab for histology, blood chemistry, and hematology assays after 24 h of recovery as detailed below in “Tissue collection and histopathology”. The same exact process was done on another 4 male and 4 female mice using phosphate buffered saline (PBS) as a control. The same process was repeated on another 16 mice (8 male and 8 female, evenly divided between the ampyrone group and the PBS control group with equal number of animals of each sex in each group), except these mice were submitted to the veterinary service after a 14-day recovery period.

### Tissue collection and histopathology

Mice were euthanized by CO_2_ asphyxiation and cardiac exsanguination. Terminal cardiac blood was collected for complete blood counts (CBC) and serum chemistry evaluation through the Stanford Animal Diagnostic Laboratory. The abdominal skin and subcutaneous tissues were collected and immersion fixed in 10% neutral buffered formalin for 72 h. Following fixation, two longitudinal sections of abdominal skin were collected along the ventral midline, parallel to hair follicle growth. Formalin-fixed skin samples were submitted to HistoTec Laboratories (Hayward, CA), processed routinely, embedded in paraffin, sectioned, and stained with hematoxylin and eosin (H&E). H&E samples were evaluated by a board-certified veterinary pathologist using an ordinal scale (scores 0-4) where scores reflect the percentage of tissue exhibiting a given finding. Scores were assigned as follows: 0 = absent; 1= minimal (<25% of sample), 2 = mild (25-50% of sample), 3 = moderate (50-75% of sample), 4 = severe (>75% of sample). Skin samples were scored for the following parameters: mucosal ulceration; follicular and dermal necrosis +/− inflammation; superficial bacteria; acanthosis and hyperkeratosis; serocellular crusts; dermal fibrosis; follicular and adnexal dropout or disorganization; lymphoplasmacytic dermatitis.

### Replication

The sample size for each experiment was determined by power analysis to ensure statistical rigor for all comparisons. Sample size, *n*, was calculated as:

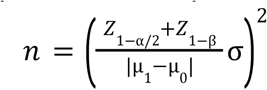

Where α=0.05, *Z*_1−α/2_ =1.96, *Z*_1−β_ =0.84, β = 0. 20, µ_1_ is the mean of the outcome variable for the experimental condition, µ_0_ is the mean of the outcome variable for the control, and σ is the standard deviation of the outcome variable.

### Statistics

Comparison between groups was evaluated with a two-tailed, heteroscedastic, t-test. P<0.05 was considered statistically significant.

### Data, Materials, and Software Availability

All data are available in the research article or in the SI Appendix. Solutions and hydrogels containing absorbing molecules are available from G.H. under a materials transfer agreement with Stanford University.

## Acknowledgements

Ellipsometry characterizations were performed at the Stanford Nano Shared Facilities (SNSF), supported by the National Science Foundation under award ECCS-1542152. Any opinions, findings, and conclusions, or recommendations expressed in this material are those of the authors and do not necessarily reflect the views of the National Science Foundation. *In vivo* two-photon imaging experiments were performed at Stanford Wu Tsai Neuroscience Microscopy Service (NMS). HPLC-MS data for ampyrone was collected by the Stanford University Mass Spectrometry (SUMS) core facility. ^1^H NMR was done at Stanford Sarafan ChEM-H Medicinal Chemistry Knowledge Center (MCKC). We gratefully acknowledge Z. Ou for useful discussions and assistance.

## Funding

G.H. acknowledges three awards from the NIH (5R00AG056636-04, 1R34NS127103-01, and R01NS126076), an NSF CAREER award (2045120), an NSF EAGER award (2217582), a Rita Allen Foundation Scholars Award, a Beckman Technology Development Grant, a grant from the Focused Ultrasound Foundation, a gift from the Spinal Muscular Atrophy (SMA) Foundation, gifts from the Pinetops Foundation, two seed grants from the Wu Tsai Neurosciences Institute, two seed grants from the Bio-X Initiative of Stanford University, and a teacher-scholar award from the Camille and Henry Dreyfus Foundation. M.L.B. acknowledges a grant from the Air Force Office of Scientific Research (FA9550-21-1-0312). C.H.C.K. acknowledges the National Science Foundation Graduate Research Fellowships program (1656518) and the Wu Tsai Neuroscience NeuroTech Training program. E.L.S. acknowledges a TIME fellowship. R.H.R. acknowledges NIH/NINDS K99NS130078. B.M.F. acknowledges a Howard Hughes Medical Institute funded fellowship through the Damon Runyon Cancer Research Foundation (DRG 2518-24). C.R.B. acknowledges an award from the NIH (R01CA200423).

## Author Contributions

Conceptualization: C.H.C.K., E.L.S., and G.H.

Dye screening: C.H.C.K., E.L.S., and S.Z.

Materials absorption characterization: C.H.C.K., E.L.S., and A.P.

Ellipsometry characterization: C.H.C.K. and A.P.

Silica phantom characterization: C.H.C.K., E.L.S., and A.P.

*Ex vivo* tissue characterization: C.H.C.K.

Brightfield mouse imaging: C.H.C.K.

Hydrogel fabrication: M.C., X.C., and C.W.

Mouse breeding: C.H.C.K., R.H.R., S.C., L.Z., Q.Y., L.Y., and A.B.

AAV injection: R.H.R. and C.H.C.K.

Two photon excitation fluorescence microscopy: C.H.C.K., R.H.R., G.X.W., and E.L.S.

Skin protein extraction: E.L.S., A.T., E.R.L., and C.H.C.K.

Quantitative proteomics: E.L.S. and B.M.F.

Video motion correction: H.B.G. and T.P.C.

Histology and blood assays: K.M.C. and W.R.

Equipment fabrication: C.H.C.K.

Technical assistance: D.E., H.C., L.C., and K.B.

Supervision: M.L.B., J.B.D., and G.H.

Original draft: C.H.C.K., E.L.S., B.M.F., and G.H.

Review and editing: C.H.C.K., E.L.S., P.A.L., M.L.B., J.B.D., and G.H.

## Competing Interest Statement

No competing interest

## Notes

### Competing Interest Statement

The authors have declared no competing interest.

