## Supporting Information for "Color-neutral and reversible tissue transparency enables longitudinal deep-tissue imaging in live mice"

Color-neutral and reversible tissue transparency enables longitudinal deep-tissue imaging in the same animals

**This PDF file includes:**

Supporting text
Figures S1 to S8
Tables S1 to S2
Movies S1 to S2
Datasets S1 to S2
Supporting references

### Supporting Text

**Optimal age range of mice for through-scalp imaging**. Precisely controlling the exact age of the mice is essential for achieving optimal scalp transparency. We have found that in mice younger than P15, the skull itself severely aberrates the 920 nm excitation light required for two-photon microscopy, despite appearing visibly transparent. Since two-photon microscopy is more sensitive to excitation aberrations than emission aberrations, imaging through the skull of P14 and younger mice becomes challenging. After approximately P28, the skull becomes too thick, introducing severe aberrations that prevent cortical imaging. Because our approach does not render the skull transparent, achieving optical transparency in the scalp alone is insufficient for microscopic imaging of the cortex through both the scalp and skull, if the skull causes significant aberrations. Therefore, P14–P28 represents the optimal age range for through-scalp two-photon microscopy using our scalp transparency approach.

**Recovery from anesthesia and ampyrone.** Ampyrone is an analgesic (1), so careful tuning of anesthetic dosage is crucial during animal experiments involving ampyrone. To minimize combined exposure, we recommend using inhalational (e.g., isoflurane) rather than intraperitoneal anesthesia (e.g., ketamine), as the former allows real-time dose adjustments during procedures. Isoflurane dosing and flow rate should be continuously monitored and adjusted based on visual assessment of the mouse’s heart and breathing rates.

For optimal workflow, one researcher should focus on skin treatment and imaging, while another manages anesthesia. Prolonged exposure to both ampyrone and anesthesia can hinder recovery. However, when ampyrone is administered without anesthesia for awake experiments, mice can tolerate over an hour of exposure without recovery issues (Movie S1).

That said, prolonged ampyrone application to the skin, even in awake mice, reduces the likelihood of full skin recovery. To optimize recovery, it is best to keep the skin transparency area to approximately 1 $cm^2$ and duration to approximately 20-40 min.

**Transparency effect in the mouse scalp.** Unlike with the abdomen, reversing scalp transparency for longitudinal two-photon microscopy presented challenges, as we were unable to apply a bioadhesive hydrogel patch and secure it with surgical tape. This difficulty arises from the inability to attach any foreign object to a mouse’s head without the animal eventually removing it. Unlike chronic brain implants, which are directly cemented to the skull, our approach preserves the scalp, making it an unstable platform for securing external materials.

As a result, we applied Eucerin Advanced Repair Cream (NART #87955, Eucerin, Beiersdorf AG, Germany) to the scalp every 30 minutes for the first 5 hours, then every 8 hours thereafter. However, this treatment is less effective than the bioadhesive hydrogel, and the scalp remains partly translucent for several days following ampyrone treatment, reversal via thorough PBS rinsing, and Eucerin application.

**Interpretation of histology results.** After 1 day after treatment, mice in the control (PBS) and experimental (ampyrone) group exhibited similar levels of ulceration and dermal necrosis. At 14 days after treatment, mice in both groups exhibited similar levels of dermal fibrosis and follicular and adnexal drop out. It should be noted that while only the removal of the stratum corneum is necessary for the diffusion of ampyrone through the skin, these histological images show that some deeper layers of the epidermis are also partially disrupted. This inadvertent disruption results from excessive mechanical rubbing of the skin, which can be minimized or avoided by applying gentler force. Furthermore, the fact that skin treated with PBS shows similar or even greater levels of epidermal disruption than ampyrone suggests that ampyrone itself did not cause this damage.

### Supplementary Figures

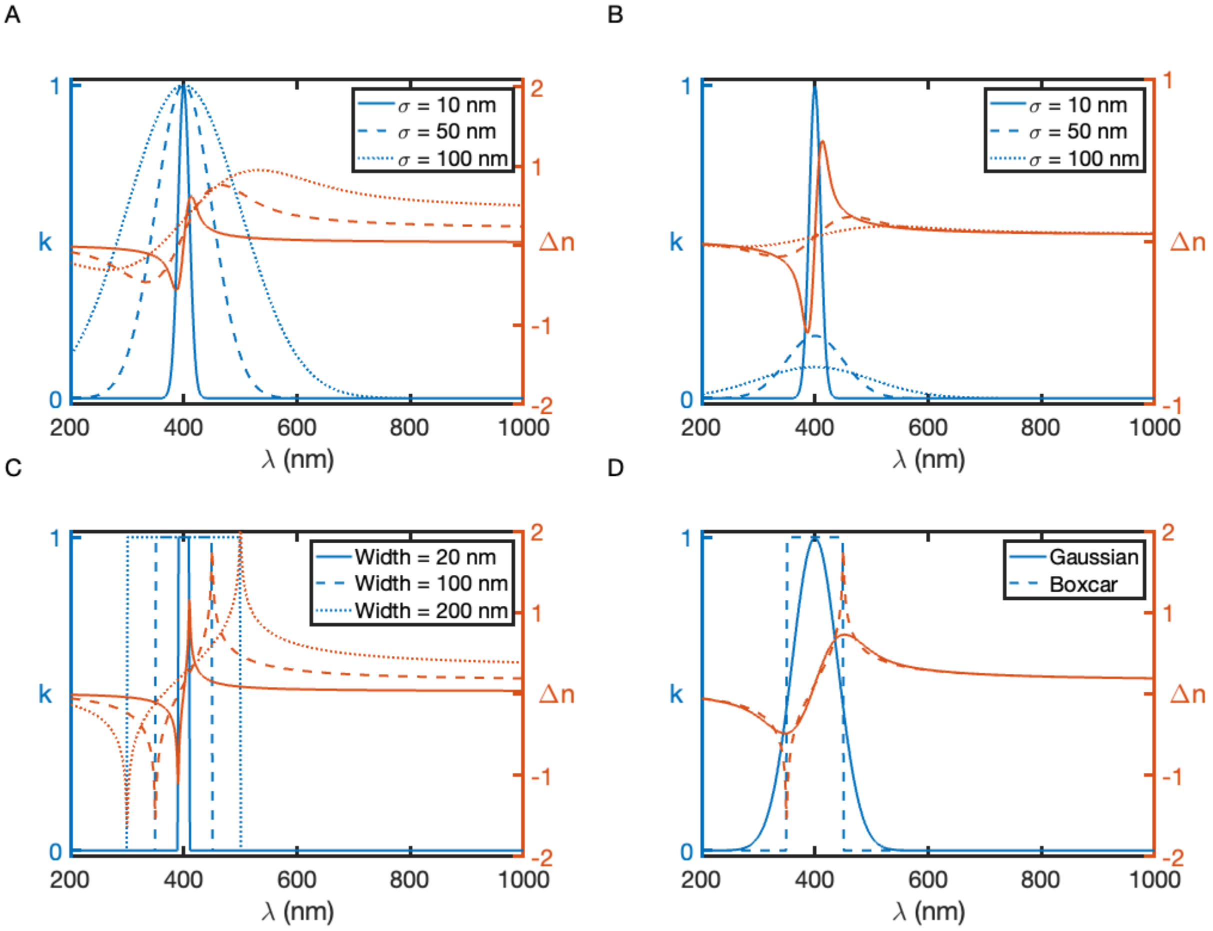


**Fig. S1.** Kramers-Kronig simulations of absorption spectra modeled as Gaussian and boxcar functions. (*A*) Absorption spectra modeled as Gaussian functions (blue) with the corresponding RI modulation (orange). The standard deviation of the Gaussian function, σ, is set to 10, 50, and 100 nm with the height kept constant. (*B*) Absorption spectra modeled as Gaussian functions (blue) with the corresponding RI modulation (orange). The standard deviation of the Gaussian function, σ, is set to 10, 50, and 100 nm with the area under the curve kept constant. (*C*) Absorption spectra modeled as boxcar functions (blue) with the corresponding RI modulation (orange). The width of the boxcar function is set to 20, 100, and 200 nm with the height kept constant. (*D*) Absorption spectra modeled as Gaussian and boxcar functions with the same area under the curve (blue) alongside the corresponding RI modulation (orange). These plots, taken together, show that far-off-resonance RI modulation depends on total area under the curve of the absorption spectrum, while near-resonance depends on the sharpness of the absorption peak.

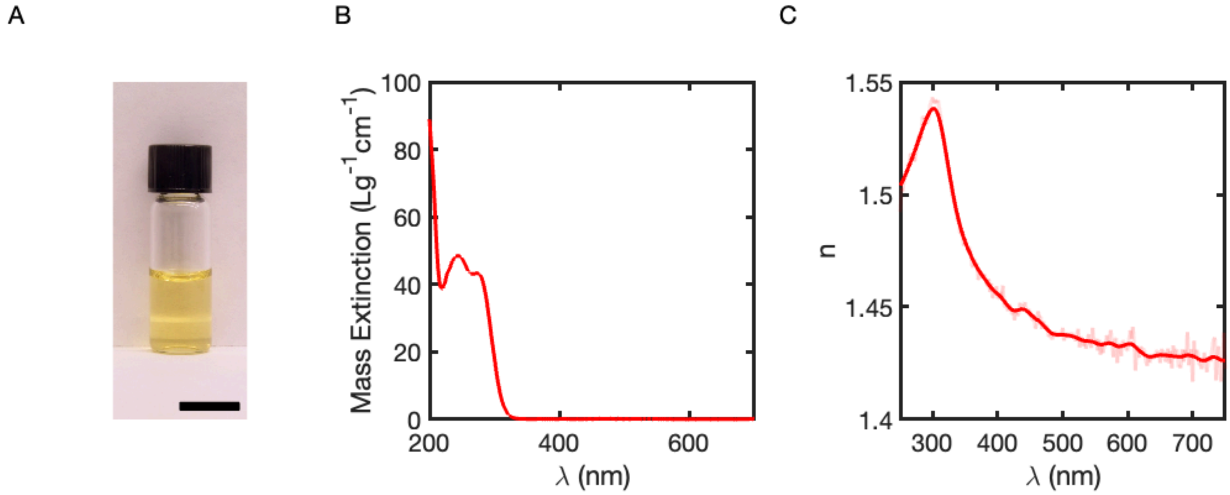


**Fig. S2.** (*A*) 38% w/w ampyrone dissolved in DI water. Scale bar: 1 cm. (*B*) Full-range mass extinction spectrum of 0.01% w/w ampyrone in DI water with 1 mm pathlength cuvette. (*C*) Bulk refractive index spectrum of 38% w/w ampyrone dissolved in DI water.

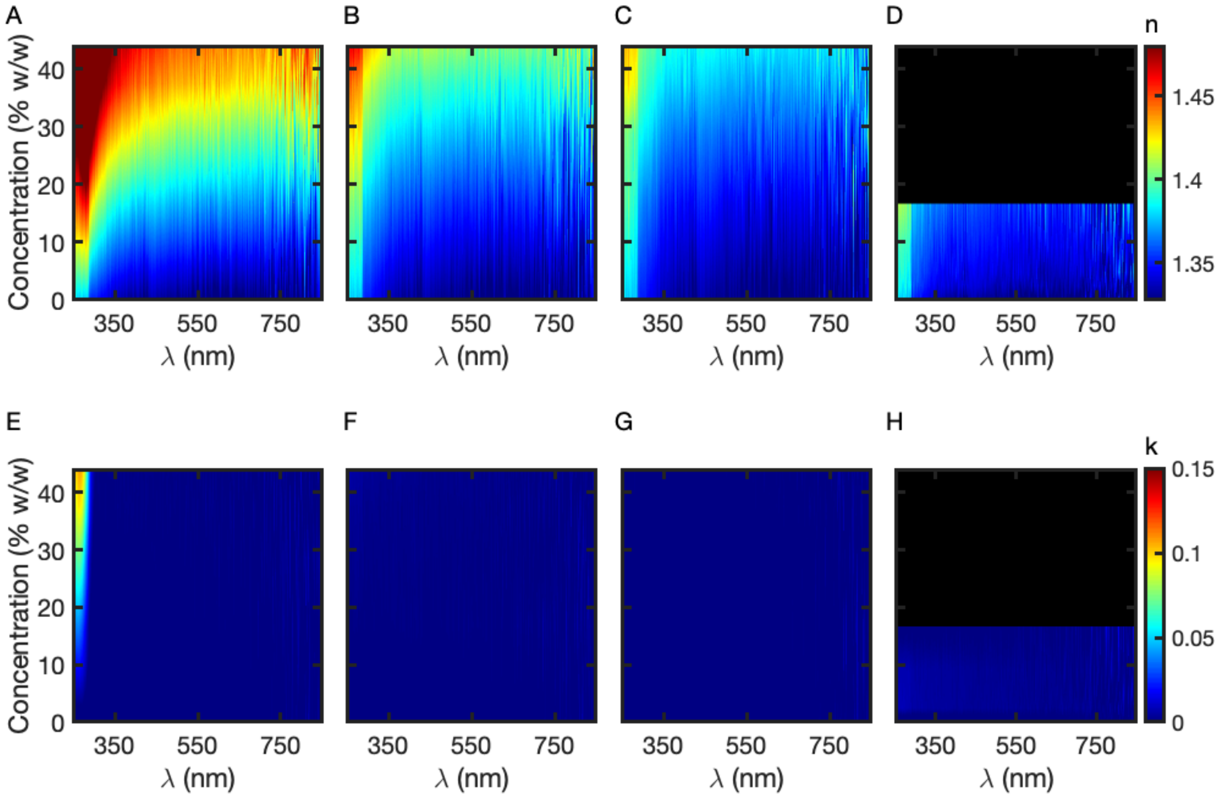


**Fig. S3.** Ellipsometry data for all compounds in Fig. 1. Real RI spectra of (*A*) phenazone, (*B*) sucrose, (*C*) glycerol, and (*D*) dextran. Imaginary RI spectra of (*E*) phenazone, (*F*) sucrose, (*G*) glycerol, and (*H*) dextran. Black shading in the plots (*D&H*) indicates unavailable data due to solubility constraints.

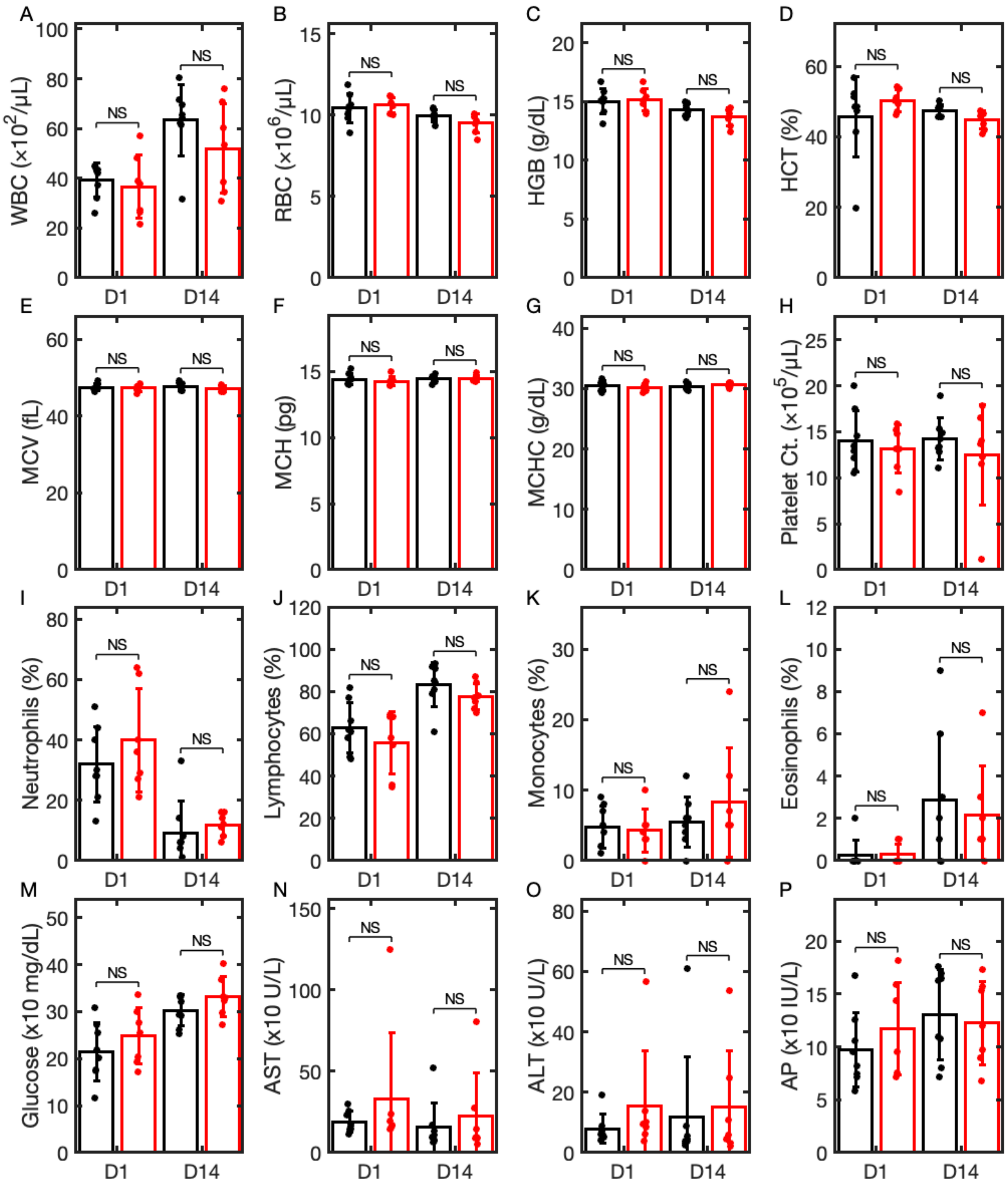


**Fig. S4.** Blood panel results. Mice in the experimental group were topically treated with 38% w/w ampyrone (red) while mice in the control group were treated with 1x PBS in an identical manner (black). All mice were allowed to recover using a topically applied hydrogel. D1 mice were collected after 1 day of recovery from topical application of ampyrone or PBS. D14 mice were collected after 14 days of recovery. (*A*) White blood cells. (*B*) Red blood cells. (*C*) Hemoglobin. (*D*) Hematocrit. (*E*) Mean corpuscular volume. (*F*) Mean corpuscular hemoglobin. (*G*) Mean corpuscular hemoglobin concentration. (*H*) Platelet count. (*I*) Neutrophils. (*J*) Lymphocytes. (*K*) Monocytes. (*L*) Eosinophils. (*M*) Glucose. (*N*) Aspartate transaminase. (*O*) Alanine transaminase. (*P*) Alkaline phosphatase. Sample size is n=8 mice for PBS D1, n=7 for ampyrone D1, n=7 for PBS D14, and n=8 for ampyrone D14. Bar graphs are shown as mean values ± SD.

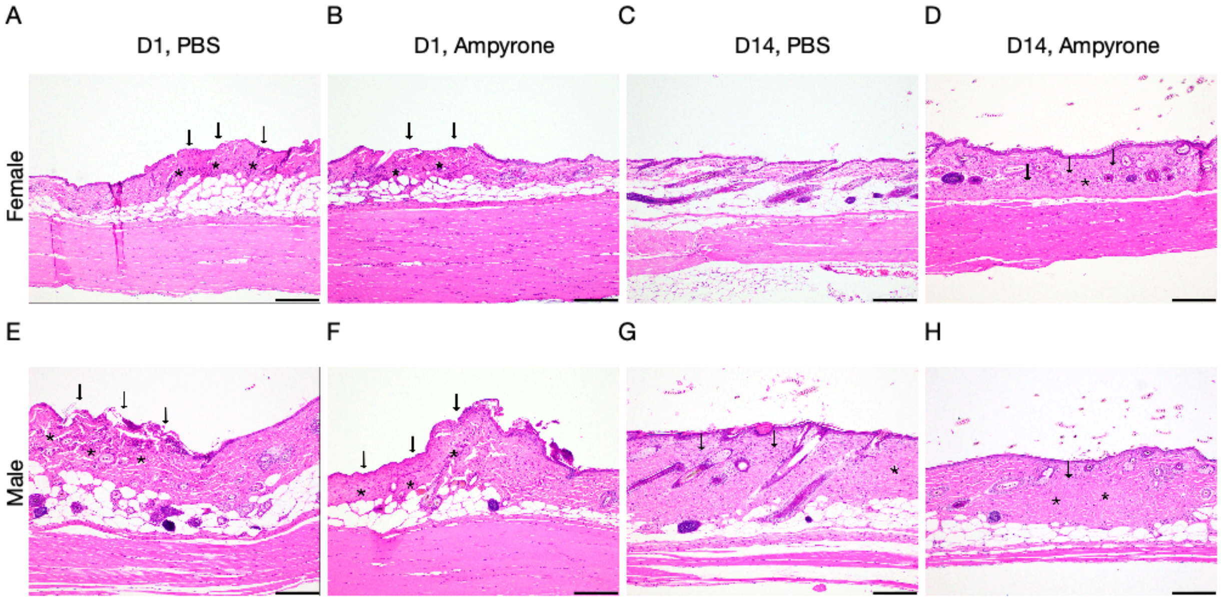


**Fig. S5.** Histology results for mice abdominal skin treated with PBS and ampyrone. (*A*-*D*) Histology images of abdominal skin from representative female mice on 1 day after treatment with PBS (*A*), 1 day after treatment with ampyrone (*B*), 14 days after treatment with PBS (*C*), and 14 days after treatment with ampyrone (*D*). (*E*-*H*) Histology images of abdominal skin from representative male mice on 1 day after treatment with PBS (*E*), 1 day after treatment with ampyrone (*F*), 14 days after treatment with PBS (*G*), and 14 days after treatment with ampyrone (*H*). For all day 1 images, arrows indicate ulceration and asterisks indicate dermal necrosis while for all day 14 images, arrows indicate dermal fibrosis and asterisks indicate follicular and adnexal drop out.Scale bars are all 200 μm.

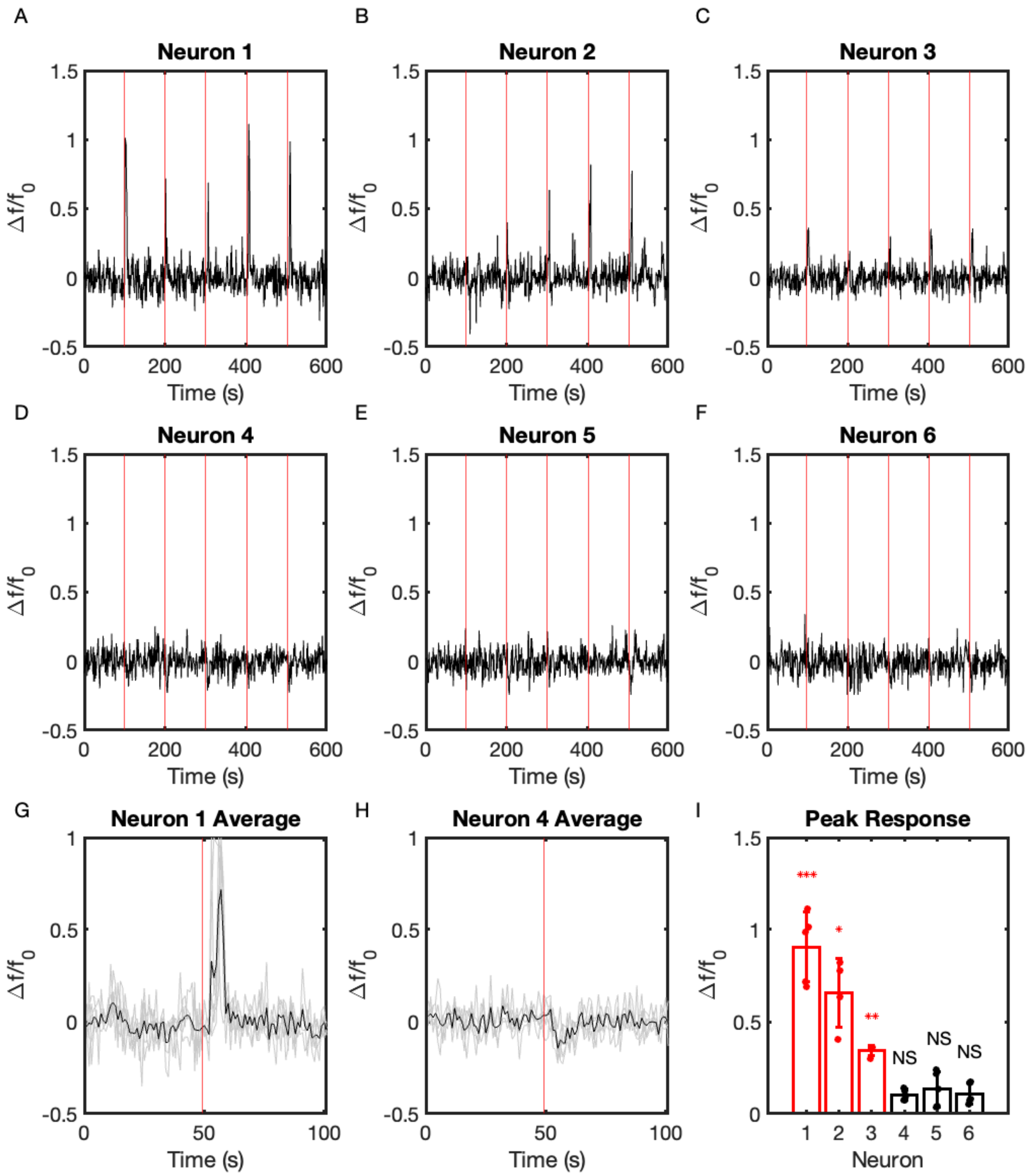


**Fig. S6.** Individual neuron traces from Fig. *6*. (*A-F)* Individual traces of the dynamic calcium signals corresponding to those in Fig. *6F*. Red lines indicate the stimulus applied to the mouse. (*G)* Average (black) of all transients for neuron 1 as an example of a responsive neuron. Individual traces are shown in gray. (*H)* Average (black) of all transients for neuron 4 as an example of a non-responsive neuron. Individual traces are shown in gray. (*I*) Peak responses for all 6 neurons, with the red data for the active neurons showing a much greater peak response than their inactive counterparts. The P-values between the peak and baseline responses for neurons 1-3 are 0.00055 (***), 0.019 (*), and 0.0022 (**). The P-values between the peak and baseline responses for neurons 4-6 are 0.058, 0.066, and 0.92 (NS, not significant).

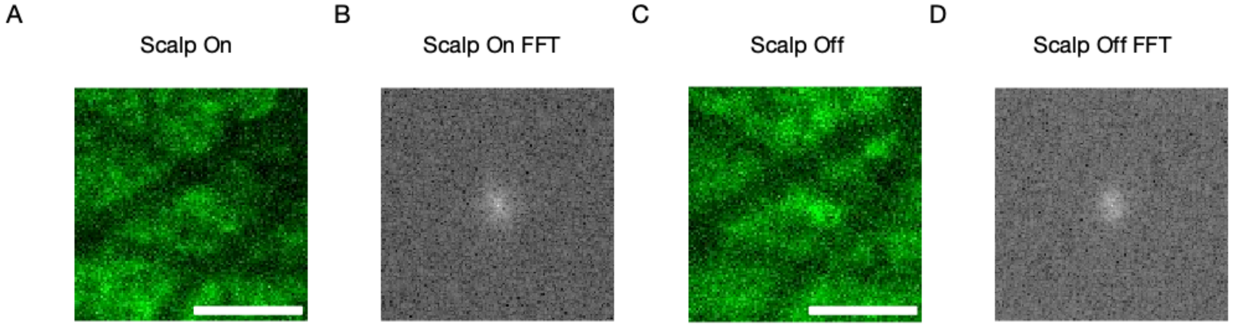


**Fig. S7.** Comparison of GCaMP images of the same cortical region in a mouse through the transparent scalp and with the scalp removed. (*A*) GCaMP image of the mouse cortex through the transparent scalp. (*B*) The corresponding fast Fourier transform (FFT) analysis for *A*. (*C*) GCaMP image of the mouse cortex with scalp removed and skull exposed. (*D*) The corresponding FFT analysis for *C*. In the FFT images, low spatial frequencies are concentrated at the center, while higher spatial frequencies increase toward the edges. The relative diameter of the central spot suggests that the resolvable higher spatial frequencies are similar with and without the scalp, indicating comparable imaging resolution in both conditions. Scale bars are 200 µm.

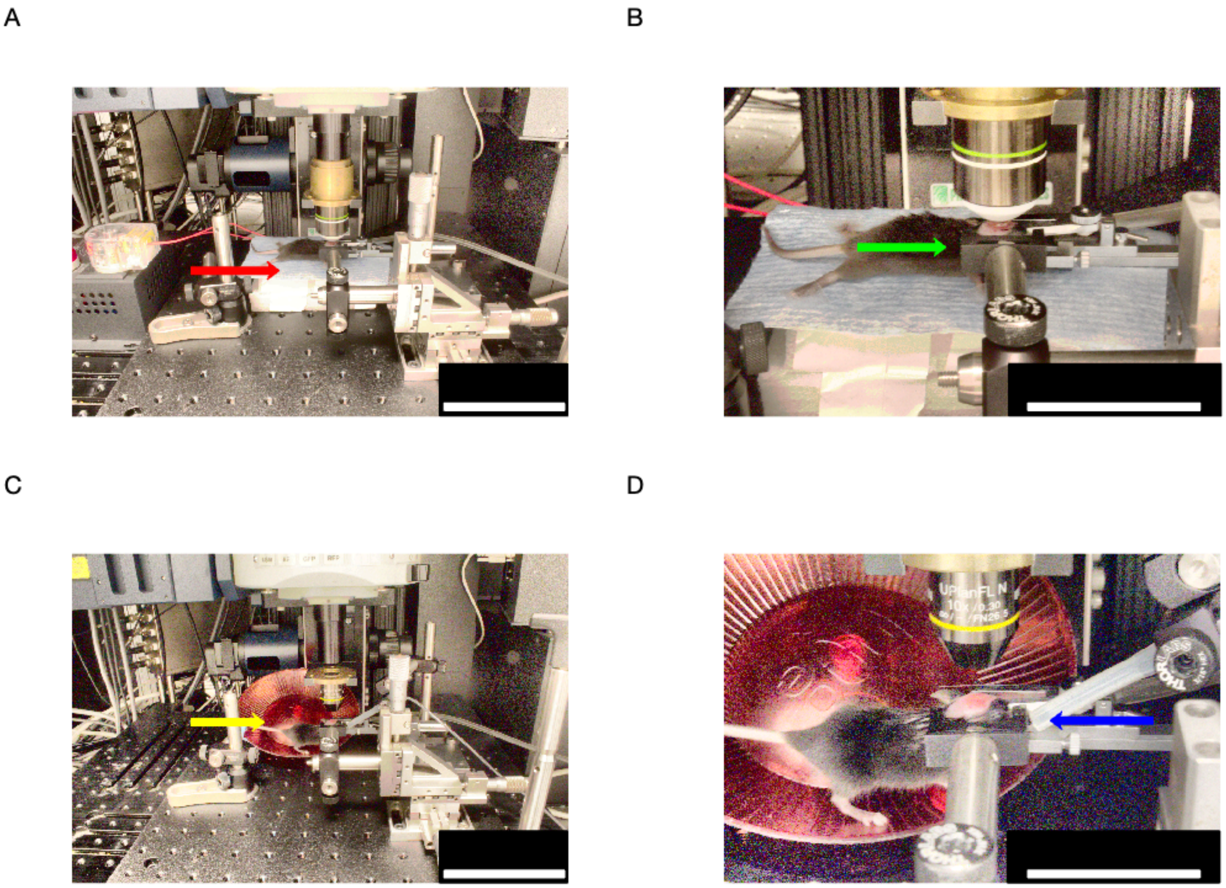

**Fig. S8.** Two-photon imaging setups for live mice. (*A*) Two-photon fluorescence microscopy setup for a mouse under anesthesia, showing the heating pad (red arrow) that maintains the mouse body temperature during imaging. (*B*) The same two-photon fluorescence microscopy setup as in *A* at higher magnification, showing the coverslip holder (green arrow) that isolates the objective from the mouse. (*C*) Two-photon fluorescence microscopy setup for an awake mouse, showing the treadmill (yellow arrow) used to minimize movement and reduce stress. (*D*) The same two-photon fluorescence microscopy setup as in *C*, showing an air tube (blue arrow) used to deliver an air-puff stimulus to the mouse whiskers and face during calcium imaging. The anesthetized setup was used for structural, non-longitudinal YFP imaging and for GCaMP imaging with a 20X immersion objective, while the awake setup was used for longitudinal YFP imaging and functional GCaMP imaging with a 10X air objective. Scale bars are 10 cm in *A&C* and 5 cm in *B&D*.

**Table S1.** P-values comparing control and experimental conditions for Fig. S4 on both D1 and D14. A two-tailed, heteroscedastic (two-sample unequal variance, unequal data size) t-test was performed for D1 and D14 to obtain these P-values. A P-value of less than 0.05 was considered statistically significant. There was no statistically significant difference between control (PBS treatment) and experimental (ampyrone treatment) groups on either Day 1 (D1) or Day 14 (D14) based on this criterion. The acronyms correspond to WBC: white blood cells; RBC: red blood cells; HGB: hemoglobin; HCT: hematocrit; MCV: mean corpuscular volume; MCH: mean corpuscular hemoglobin; MCHC: mean corpuscular hemoglobin concentration; AST: aspartate transaminase; ALT: alanine transaminase.

| | | P-values | |
|---|---|---|---|
| | Test | **Day 1** | **Day 14** |
| Complete blood count | **WBC** | 0.65 | 0.20 |
| | **RBC** | 0.62 | 0.13 |
| | **HGB** | 0.83 | 0.11 |
| | **HCT** | 0.31 | 0.06 |
| | **MCV** | 0.81 | 0.28 |
| | **MCH** | 0.45 | 0.89 |
| | **MCHC** | 0.49 | 0.18 |
| | **Platelet Count** | 0.58 | 0.44 |
| | **Neutrophils** | 0.33 | 0.51 |
| | **Lymphocytes** | 0.32 | 0.22 |
| | **Monocytes** | 0.77 | 0.41 |
| | **Eosinophils** | 0.91 | 0.61 |
| Blood chemistry tests | **Glucose** | 0.32 | 0.16 |
| | **AST** | 0.40 | 0.57 |
| | **ALT** | 0.33 | 0.75 |
| | **Alkaline Phosphatase** | 0.36 | 0.72 |

**Table S2.** Mean skin histology rankings. For scoring, 0 = absent, 1 = minimal (<25% of sample), 2 = mild (25-50% of sample), 3 = moderate (50-75% of sample), and 4 = severe (>75% of sample). Lower scores indicate better skin recovery. No statistically significant difference is found between ampyrone-treated and PBS-treated groups on both days. All data are reported as mean ± standard deviation, SD.

| | **Weight (g)** | **Mucosal ulceration** | **Follicular and dermal necrosis** | **Superficial bacteria** | **Acanthosis and hyperkeratosis** | **Serocellular crusts** |
|---|---|---|---|---|---|---|
| **Day 1 Ampyrone** | 23±5 | 2.3±1.3 | 1.8±0.5 | 0±0 | 0.5±0.6 | 0.8±0.5 |
| **Day 1 PBS** | 22±4 | 2.8±0.5 | 2.8±0.5 | 0±0 | 0.8±0.5 | 1±0 |
| **Day 1 P-values** | 0.84 | 0.50 | 0.03 | - | 0.54 | 0.39 |
| | **Weight (g)** | **Dermal Fibrosis** | **Follicular and adnexal dropout or disorganization** | **Lymphoplasmacytic dermatitis** | **Acanthosis and hyperkeratosis** | **Serocellular crusts** |
| **Day 14 Ampyrone** | 27±5 | 1.3±0.6 | 1±0 | 1±0 | 0.7±0.6 | 0±0 |
| **Day 14 PBS** | 26±3 | 1.3±1.0 | 1.3±1.0 | 1.3±1.0 | 1.0±0.8 | 0±0 |
| **Day 14 P-values** | 0.85 | 0.89 | 0.64 | 0.64 | 0.55 | - |

**Movie S1 (**‘Movies_S1.mp4’**)** A head-fixed, awake mouse actively running on a treadmill with a transparent scalp following ampyrone treatment.

**Movie S2 (**‘Movie_S2.mp4’**)** A representative two-photon fluorescence video depicting neuronal activity via GCaMP under stimulation. FOV is 1.170 x 1.170 μm and frame rate is the native 0.987 FPS.

**Dataset S1 (**‘Dataset_S1.stl’**)** STL file for 3D printing a custom nose cone designed for delivering anesthesia to the Narishige head fixation unit. While Narishige offers a commercially available nose cone system, it lacks a nose clamp, which may result in less secure fixation, particularly for awake mice. All 3D printing was performed using a Formlabs Form 3+ with black resin, followed by ambient light and air curing to ensure dimensional accuracy.

**Dataset S2 (**‘Dataset_S2.stl’**)** STL file for 3D printing a custom coverslip holder designed to provide a flat and clean surface for imaging under two-photon excitation. The coverslip enhances stability by further restricting head motion and serves as a barrier to isolate the water immersion objective from ampyrone on the scalp surface. The coverslip holder was also used with the air objective, primarily for the aforementioned stability it provided.
